# A latent pool of neurons silenced by sensory-evoked inhibition can be recruited to enhance perception

**DOI:** 10.1101/2024.02.12.579847

**Authors:** Oliver M. Gauld, Adam M. Packer, Lloyd E. Russell, Henry W.P. Dalgleish, Maya Iuga, Francisco Sacadura, Arnd Roth, Beverley A. Clark, Michael Häusser

## Abstract

Which patterns of neural activity in sensory cortex are relevant for perceptual decision-making? To address this question, we used simultaneous two-photon calcium imaging and targeted two-photon optogenetics to probe barrel cortex activity during a perceptual discrimination task. Head-fixed mice discriminated bilateral whisker deflections and reported decisions by licking left or right. Two-photon calcium imaging revealed sparse coding of contralateral and ipsilateral whisker input in layer 2/3 while most neurons did not show task-related activity. Activating small groups of pyramidal neurons using two-photon holographic photostimulation evoked a perceptual bias that scaled with the number of neurons photostimulated. This effect was dominated by the optogenetic activation of a small number of non-coding neurons, which did not show sensory or motor-related activity during task performance. Patterned photostimulation also revealed potent recruitment of cortical inhibition during sensory processing, which strongly and preferentially suppressed non-coding neurons. Our results provide a novel perspective on the circuit basis for the sparse coding model of somatosensory processing in which a pool of non-coding neurons, selectively suppressed by strong network inhibition during whisker stimulation, can be recruited to enhance perception.

**Highlights:**

- All-optical interrogation of barrel cortex during bilateral whisker discrimination
- Sparse coding of contralateral and ipsilateral whisker information
- Selective sensory-evoked inhibition helps ensure sparse coding
- Optogenetic recruitment of stimulus non-coding neurons can aid perception

## INTRODUCTION

Understanding how sensory inputs are transformed into perceptual outputs requires functional dissection of cortical circuits during behavior^1,2^. Neural circuits in superficial layers of sensory cortex are largely composed of excitatory neurons that show heterogenous stimulus tuning^3^, structured patterns of connectivity^4–6^, and low spike rates^7^. GABAergic interneurons, which are densely connected with local excitatory neurons^8,9^ and have high baseline firing rates^10^ and broad receptive fields^11^, provide inhibition that patterns spatiotemporal excitatory dynamics^12–16^. Accordingly, sensory processing in superficial cortex is typically dominated by subsets of highly-tuned neurons^17–19^, with most excitatory neurons remaining ‘silent’ during behavior^20,21^.

The observation that few neurons are engaged during sensory processing, and that their activity correlates with perceptual decisions^22–24^, suggests cortex uses a sparse neural code to generate stimulus percepts^23,25–27^. Sparse coding is an efficient mechanism for encoding information^28–30^, and has been observed experimentally across a range of neural systems and species^31,32^. In causal support of the sparse coding hypothesis, experimental stimulation of small groups of cortical neurons, and even single neurons, can elicit perceptual responses^24,33–39^. Moreover, functionally targeted cortical microstimulation can also influence decisions in favour of the tuning of the manipulated neurons^40–45^.

As these findings indicate that perception is driven by small stimulus-tuned ensembles, the functional significance of the large proportion of non-responsive neurons in sensory cortex has remained enigmatic^21,46,47^. On the one hand, neurons may appear silent if they have highly selective receptive fields that are not explored by standard experimental sensory stimulation paradigms^48^. Alternatively, non-responsive neurons may be reserved for implementing circuit plasticity^20,49^. Understanding why few neurons respond strongly to sensory input, and how such sparse activity can drive reliable sensorimotor behavior, is a fundamental goal for understanding brain function^19,50–52^.

Here, we used a combination of experimental approaches, including simultaneous two-photon calcium imaging and holographic two-photon photostimulation^53–55^, to interrogate sparse coding in barrel cortex while head-fixed mice performed a bilateral whisker discrimination task. We opted to probe barrel cortex under bilateral sensory conditions as bilateral processing is an important and ethological aspect of tactile sensation, especially for rodents locomoting in complex subterranean habitats^56–59^. Moreover, lesioning^56^ and pharmacologically silencing^58,60^ barrel cortex impairs bilateral whisker behavior, suggesting that bilateral somatosensation is an ethological and barrel cortex-dependent perceptual function.

First, we used large-scale optogenetic manipulations to confirm that unilateral modulation of barrel cortex activity influences contralateral whisker perception. Population calcium imaging revealed sparse coding of both contralateral and ipsilateral whisker input in L2/3, while most neurons were ‘non-coding’, as they did not show task-related activity. We then characterized the L2/3 circuit response to paired whisker stimulation and patterned two-photon photostimulation, while simultaneously monitoring ongoing changes in perceptual choices. Surprisingly, targeted two-photon photostimulation of small neuronal ensembles evoked lateralized choice biases that scaled with the number of stimulus non-coding neurons activated by targeted photostimulation. Our findings also demonstrate that strong and selective inhibitory pressure suppresses weak and non-coding neurons in the circuit, enforcing sparse circuit activity during whisker processing. This provides evidence that intrahemispheric perceptual signals can be enhanced by releasing non-coding L2/3 neurons from inhibition, consistent with work implicating ‘silent’ cortical neurons in sensorimotor plasticity^20,49^.

## RESULTS

### A bilateral discrimination task for probing whisker perception

To probe the relationship between neural coding in barrel cortex and stimulus perception we developed a bilateral whisker discrimination task for head-fixed mice. First, we expressed the calcium indicator GCaMP6s and excitatory opsin C1V1 in barrel cortex neurons using a viral expression strategy (Figure S1A & S1B). We implanted mice with a headplate and cranial window and trimmed the surrounding whiskers to isolate the C2 whiskers on either side of the snout. During the task, the contralateral and ipsilateral whiskers were threaded into glass capillaries and simultaneously deflected (Figure 1A). Mice discriminated the larger amplitude side and reported their choice with directional licking at a dual lickport to obtain water rewards (Figure 1B & 1C). We counterbalanced the stimulus-response contingency across mice (Figure 1D). Mice trained on the symmetric contingency learned a congruent spatial mapping between the stronger stimulus side and the target lick response (e.g. Stim left → Lick left), while mice trained on the asymmetric contingency learned the inverse rule (e.g. Stim left → Lick right).

**Figure 1.**
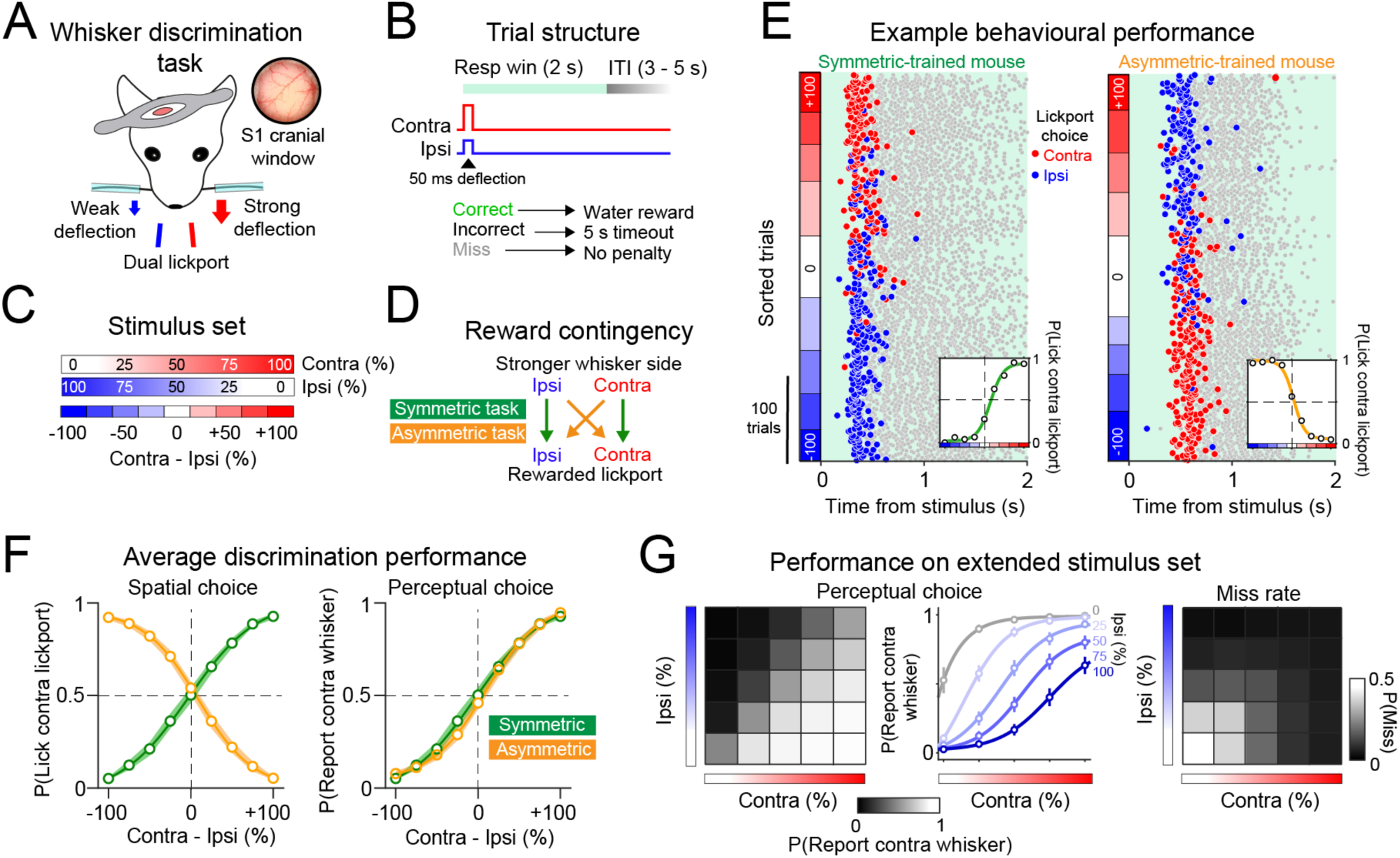
A whisker-guided task for graded bilateral intensity discrimination. **(A)** Whisker discrimination task setup. Head-fixed mice received bilateral deflections to the C2 whiskers and reported choices with directional licking. The inset shows an image of cortex viewed through a 3 mm cranial window implanted over S1. **(B)** Schematic of trial structure and trial outcome. **(C)** ‘Cross-fading’ bilateral whisker deflection stimulus design. **(D)** Overview of symmetric (green) and asymmetric (orange) stimulus-reward contingencies. **(E)** Example discrimination session performance from a symmetric-trained mouse (left) and an asymmetric-trained mouse (right). Trials were delivered in a randomised order but are sorted along the y-axis according to stimulus difference as in (C). Each row corresponds to a single trial, and each marker corresponds to a lick. The first lick response is coloured red or blue for contra or ipsi lickport choice. The inset shows the session psychometric curve. **(F)** Average psychometric performance for symmetric (green; n = 31 mice) and asymmetric (orange; n = 30 mice) trained mice. The left and right plots show discrimination performance plotted as spatial choice ‘P(Lick contra lickport)’ and perceptual choice ‘P(Report contra whisker)’ respectively. **(G)** Average performance during extended stimulus set training sessions. Left: Average P(Report contra whisker) is shown across trial-types with each square in the 5 × 5 grid representing a different combination of contra and ipsi input. Middle: Behavioural data are replotted such that each row in the left behavioural matrix (corresponding to a different ipsi stimulus level) is now shown as a psychometric curve. Right: Average probability of missing a trial is shown across stimulus conditions (n = 83 sessions in 21 mice). Group data in Figure 1 are shown as the mean across mice, with error bars representing SEM.

Mice learned the task structure through daily training on unilateral contra and ipsi trials (Figure S2A & S2B) and were considered expert when performance on both stimulus-sides reached 70% on 3 consecutive days (days to reach expert: sym 6.1 ± 3.7; mean ± std; n = 31 mice, asym 6.8 ± 5.2; n = 30 mice; n.s. P > 0.05; Wilcoxon rank-sum test). Performance was comparable across sides (Figure S2C) and contingencies (Figure S2D), although reaction times (RT) were faster for symmetric-trained mice (RT sym 448.2 ± 123.1 ms; mean ± std; asym 526.1 ± 220.3 ms, * P < 0.05 Wilcoxon rank-sum test; Figure S2E). When using weaker stimuli, miss rate and RT increased (Figure S2F & S2G) and unilateral whisker-trimming selectively abolished detection on the trimmed (P(Miss) *baseline* 0.02 ± 0.03; *trimmed whisker* 0.84 ± 0.19 mean ± std, *** P < 0.0001, Wilcoxon signed-rank test; n = 8 mice), but not the spared (P(Miss) *baseline* 0.04 ± 0.04, *spared whisker* 0.04 ± 0.07, n.s. P > 0.05), whisker side (Figure S2H). Muscimol infusion into barrel cortex impaired detection of contralateral trials (P(Miss) *contra whisker baseline*: 0.02 ± 0.04, +*muscimol* 0.64 ± 0.2 mean ± std, ** P < 0.01 Wilcoxon signed-rank test, n = 4 mice), but not ipsilateral trials (P(Miss) *ipsi whisker baseline* 0.02 ± 0.03, +*muscimol* 0.18 ± 0.16, n.s. P > 0.05; Figure S2I), indicating that stimulus detection requires both whisker input and contralateral barrel cortex.

Once mice reached expert performance, we transitioned them to bilateral discrimination training. Single-session performance yielded high-quality psychometric curves with large numbers of trials per session (309.1 ± 93.7 trials; mean ± std; Figure 1E). Average psychometric curves from symmetric and asymmetric-trained cohorts of mice were inverted when we quantified spatial choice tendency as a function of stimulus difference (Figure 1F *left; P(Lick contra lickport)*), but comparable when we quantified the tendency mice would report the contralateral whisker stimulus (Figure 1F *right*; *P(Report contra whisker)*). As the comparison between contingencies was not the primary focus of our study, we subsequently pooled data across task contingencies unless otherwise stated. We trained some mice on an extended stimulus set comprising a larger combination of trial-types (5 × 5 stimulus ‘matrix’; n = 83 sessions in 21 mice). Discrimination performance during these sessions demonstrates that mice solve the task by integrating stimuli bilaterally, as opposed to relying on a unilateral strategy (Figure 1G). We also probed the temporal limits of bilateral discrimination (Figure S3A; n = 58 sessions in 25 mice). Temporal intervals in bilateral stimulation led to strong choice biases that aligned with the leading stimulus side and saturated at 100 ms (Figure S3B & S3C). Bilateral temporal sensitivity appeared stronger in symmetric-trained mice (Figure S3D) and correlated with average reaction times (RT) on unilateral whisker trials (Figure S3E). Together, our results demonstrate that the whisker system supports fine-scale discrimination of bilateral tactile input and can rapidly transform stimulus percepts into goal-directed motor output.

### Bulk optogenetic activation of barrel cortex evokes a contralateral percept

Following learning, we used optogenetics to probe the sufficiency of barrel cortex for whisker deflection perception. First, we photostimulated C1V1-expressing pyramidal neurons using an LED and measured the tendency for a single LED-pulse to ‘fool’ mice into reporting a contralateral whisker deflection (Figure 2A). We calculated a ‘fooling index’ as the difference in probability that optogenetic stimulation would evoke a contralateral vs an ipsilateral whisker percept as indicated by directional licking. Bulk stimulation of barrel cortex evoked illusory perceptual responses that increased with stimulation power (Fooling index: 0 mW = 0.02 ± 0.04; 10 mW = 0.11 ± 0.16; * P = 0.047; 30 mW = 0.23 ± 0.2; ** P = 0.002; 50 mW = 0.21 ± 0.17; *** P = 6.1 × 10^−5^; Wilcoxon signed-rank test comparison with 0 mW trials; n = 16 sessions in 12 mice; Figure 2B). Optogenetic stimulation evoked contralateral lickport choices in symmetric-trained mice but ipsilateral lickport choices in asymmetric-trained mice (Figure S4A). Mean RT on whisker and LED trials were highly correlated (Pearson’s correlation (r) = 0.82, *** P = 0.0002; Figure 2C), with faster LED-evoked RT measured in symmetric-trained mice (RT on LED trials; sym = 421.1 ± 78.2 ms; asym = 555.2 ± 117.4 ms; * P = 0.03; Wilcoxon rank-sum test; Figure S4B). LED-triggered licking was absent in whisker-trained control mice that did not express C1V1 (Figure S4C). Our results therefore confirm that direct optogenetic activation of barrel cortex is sufficient to evoke a contralateral whisker percept and initiate a goal-directed action specific to the learned sensorimotor context^24,38^.

**Figure 2.**
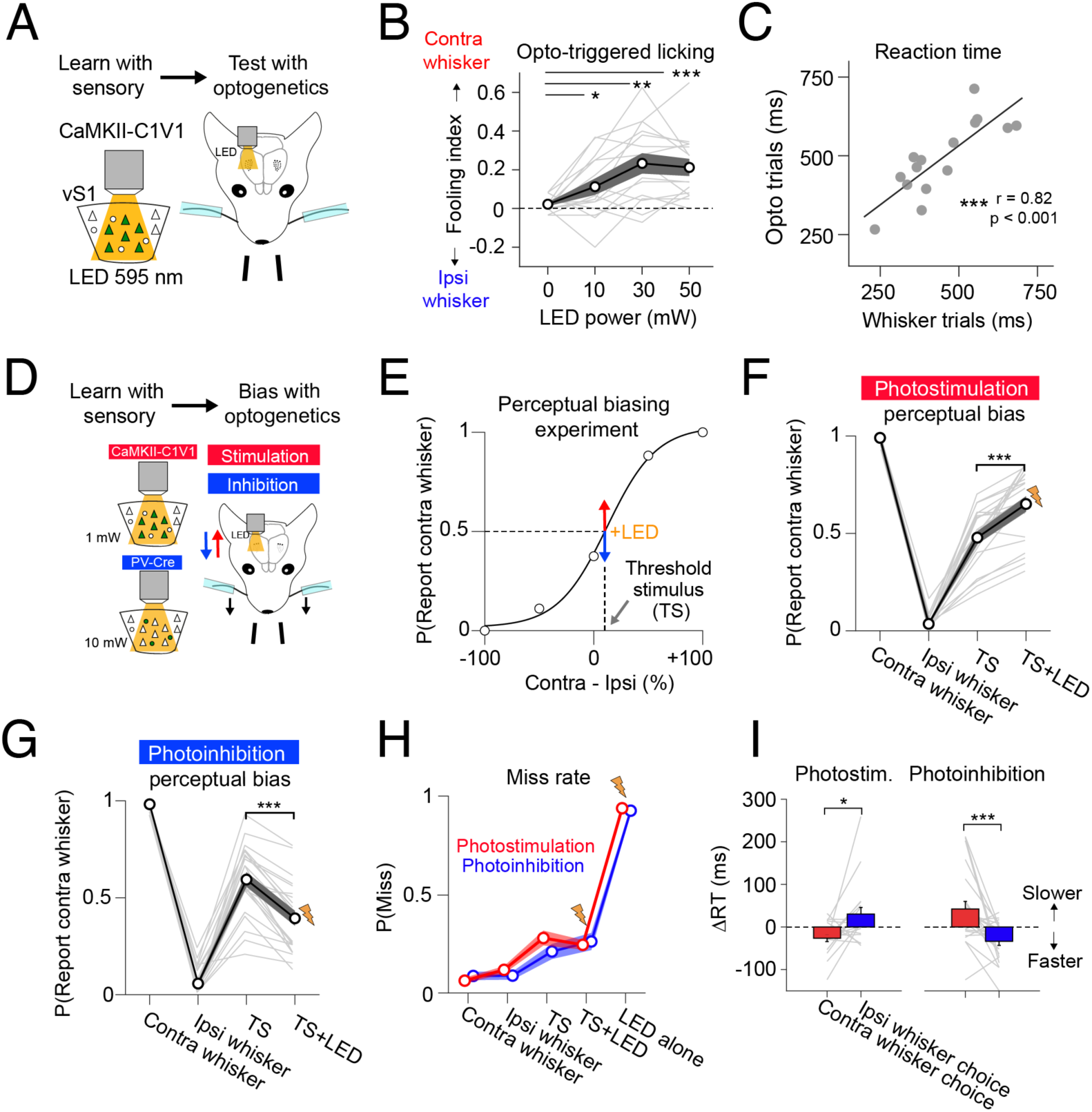
Optogenetic manipulation of barrel cortex during task-performance. **(A)** Optogenetic ‘substitution’ experiment schematic. **(B)** Perceptual ‘fooling index’ on optogenetic stimulation trials across different LED powers. Statistical comparisons were made against 0 mW ‘catch’ trials using a Wilcoxon signed-rank test. n = 16 sessions across 12 mice. **(C)** Correlation between average reaction times on optogenetic stimulation trials (mean across 30 and 50 mW trials) and whisker trials. Each data point shows an individual session (n = 16 sessions). **(D)** Schematic overview of photostimulation (red) and photoinhibition (blue) perceptual biasing experiments. Photostimulation and photoinhibition experiments were performed in 2 different sets of mice. **(E)** Optogenetic stimuli were paired with the bilateral whisker threshold stimulus (TS). **(F)** Behavioural performance during photostimulation biasing experiments. Comparison across ‘TS’ and ‘TS + PS’ trials; *** P < 0.001; Wilcoxon-signed rank test; n = 21 sessions across 19 mice. **(G)** Behavioural performance during photoinhibition biasing experiments. Comparison across ‘TS’ and ‘TS + PS’ trials; *** P < 0.001; Wilcoxon signed-rank test; n = 23 sessions across 4 mice. **(H)** Miss rate is shown across trial-types during photostimulation (red) and photoinhibition (blue) experiments. **(I)** Optogenetic biasing of TS trial reaction time for photostimulation (left) and photoinhibition (right) experiments. The mean difference in RT on trials where mice reported the contralateral whisker choice vs the ipsilateral whisker choice is shown as red and blue bars respectively. Data in Figure 2 show the average across sessions as circular markers and SEM (shaded error bars). Data from individual sessions are shown as thin grey lines. All statistical tests were two-tailed Wilcoxon signed-rank tests * P < 0.05; ** P < 0.01; *** P < 0.001.

### Bidirectional optogenetic biasing of whisker choice

Next, we assessed whether optogenetically manipulating barrel cortex biased perceptual choice at the whisker discrimination threshold (Figure 2D). We first performed photostimulation biasing experiments using a low LED power estimated to have negligible *de novo* perceptual saliency (1 mW; Figure S4D). At the start of each experiment, we calibrated the bilateral whisker threshold stimulus (TS) estimated to give rise to chance performance (Figure 2E). We then paired the TS with the LED stimulus (TS+LED trials) and varied the LED stimulus latency across trials (latency: 0 – 100 ms). To examine the perceptual effect of photostimulation we first compared average choice tendency on TS trials with all TS+LED trials. LED stimulation biased psychophysical report towards the contralateral whisker (ΔP(Report contra whisker) = 0.17 ± 0.13; *** P = 1.1 × 10^−4^; Wilcoxon signed-rank test; n = 21 sessions in 19 mice; Figure 2F), increasing contralateral lickport choices in symmetric-trained mice (ΔP(Lick contra lickport) = 0.2 ± 0.13; ** P = 0.004; Wilcoxon signed-rank test; n = 10 sessions, 8 mice) while increasing ipsilateral lickport choices in asymmetric-trained mice (ΔP(Lick contra lickport) = −0.15 ± 0.14; ** P = 0.003; Wilcoxon signed-rank test; n = 11 sessions, 11 mice; Figure S4E). The perceptual effect was strongest when LED stimulation was delivered with a short delay relative to the TS (Figure S4F), which could reflect greater temporal coincidence between optogenetic and sensory-driven activity in cortex due to the short sensory signalling latency.

We then performed experiments in a different cohort of mice to examine the perceptual effect of suppressing barrel cortex. Photostimulating C1V1-expressing parvalbumin (PV) interneurons simultaneously with TS input biased choice towards the ipsilateral whisker (ΔP(Report contra whisker) = −0.2 ± 0.11; *** P = 2.7 × 10^−5^; Wilcoxon signed-rank test; n = 23 sessions in 4 mice; Figure 2G). This effect decreased as a function of photoinhibition latency and was absent 100 ms after the TS (Figure S4G), indicating that perceptually relevant sensory processing in barrel cortex occurs within the first 100 ms from whisker stimulation. During both photostimulation and photoinhibition experiments, LED presentation did not change miss rate on TS trials (Photostimulation TS vs TS+LED ΔP(Miss) = −0.034 ± 0.13, n.s. P = 0.64; Photoinhibition ΔP(Miss) = 0.05 ± 0.13, n.s. P = 0.09) and did not evoke reliable licking responses when presented alone (P(Miss) on LED trials; photostimulation = 0.94 ± 0.06; photoinhibition = 0.93 ± 0.08; Figure 2H), indicating that perceptual effects are not mediated by strong behavioral responses to the LED stimulus alone.

Optogenetic stimulation also influenced RT on TS trials (Figure 2I). Photostimulation decreased RT for contralateral whisker choices (ΔRT contra whisker choice = −26.4 ± 36.6 ms; ** P = 0.004; Wilcoxon signed-rank test), while also tending to increase RT for ipsilateral whisker choices (ΔRT ipsi whisker choice = 30.3 ± 72.1 ms; n.s. P = 0.08; Figure 2I *left*). Photoinhibition resulted in the inverse effect, increasing RT for contralateral whisker choices (ΔRT contra whisker choice = 42.3 ± 85.2 ms; * P = 0.02), while decreasing RT for ipsilateral whisker choices (ΔRT ipsi whisker choice = −33.6 ± 45.7 ms; ** P = 0.004; Figure 2I *right*). RT effects also decreased as a function of LED-onset latency (Figure S4H). Together, our experiments show that unilateral photostimulation and photoinhibition exert inverse and time-dependent biases in perceptual decisions and perceptual speed, indicating that barrel cortex plays a causal, but transient, role in contralateral sensory processing during task performance.

### Two-photon imaging of barrel cortex during delayed-discrimination

To characterize task-related neural coding we used two-photon (2P) calcium imaging to record from GCaMP6s-expressing L2/3 pyramidal neurons. As our initial experiments indicated that mice execute rapid decisions following whisker input, we trained a new cohort of mice on a delayed-response version of the task to provide better temporal separation between ‘sensation’ and ‘action’ epochs. To circumvent the requirement to trim the surrounding whiskers, we changed the stimulus effector from a glass capillary to a ‘paddle’, which engaged multiple whiskers simultaneously (Figure 3A). To further increase stimulus saliency to aid performance in this more cognitively demanding task, the stimulus paddles were deflected using sinusoidal waveforms (duration: 500 ms) which were fixed in frequency (20 Hz) but varied in amplitude bilaterally across trials. Thus, mice solved the task by discriminating deflection amplitude bilaterally, analogous to the single-whisker task version. Following stimulus presentation, mice withheld licking across a 1 s delay until cued to respond with an auditory ‘go’ cue. A retractable lickport then moved in to initiate the response window (Figure 3B). We trained two groups of naïve mice directly on the symmetric and asymmetric version of this delayed-discrimination task (days to reach expert; sym = 8.14 ± 5; n = 7 mice; asym = 10 ± 1.6; n = 6 mice; n.s. P = 0.2; Wilcoxon ranked-sum test; Figure S5A-F). Moving the stimulus paddles out of reach of the whiskers reduced performance on unilateral trials to chance (*P(Correct) baseline* = 0.97 ± 0.03 vs *paddles out of reach* = 0.5 ± 0.06; P < 0.0001; Wilcoxon signed-rank test; n = 13 mice; Figure S5G), confirming that mice do not use audio-visual cues to solve the task. Following learning, mice were trained on the 5 × 5 ‘matrix’ stimulus-set, with discrimination performance comparable with the cohort of mice trained on the non-delay task (Figure S5H).

**Figure 3.**
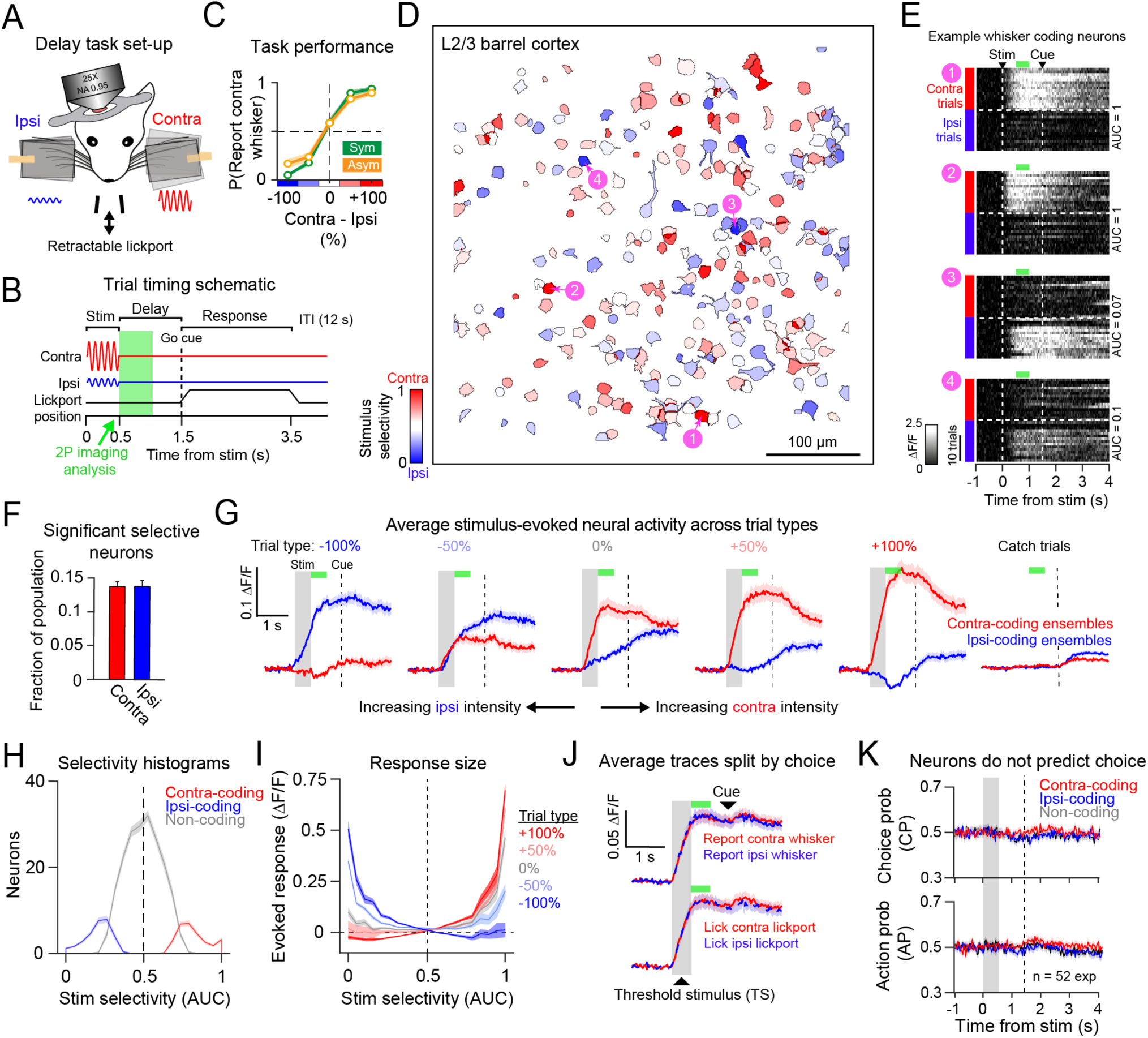
Characterisation of task-evoked activity using two-photon calcium imaging. **(A)** Schematic of the delayed-response discrimination task. **(B)** Delay-task trial structure. The green shading denotes the window used to analyse neural activity data (500 – 1000 ms). **(C)** Average psychometric performance during imaging sessions for symmetric (green; n = 7 mice) and asymmetric (orange; n = 6 mice) trained mice. **(D)** Regions of interest (ROIs) corresponding to neuronal somata in an example FOV coloured by contra vs ipsi stimulus-selectivity. **(E)** Heat-maps showing sorted single-trial fluorescence responses on unilateral contra (red) and ipsi (blue) whisker trials for 4 example neurons numbered in (D). **(F)** Quantification of the mean fraction of significant contra-coding (red) and ipsi-coding (blue) neurons per FOV (n = 52 sessions). **(G)** Average trial-evoked fluorescence traces for contra-coding neuron ensembles (red) and ipsi-coding neuronal ensembles (blue) across bilateral stimulus trial-types in (C). **(H)** Histogram showing the average distribution of stimulus-selectivity scores for contra-coding (red), ipsi-coding (blue) and stimulus non-coding (grey) neurons. **(I)** Evoked response amplitude across trial-types shown as a function of stimulus selectivity. Colours correspond to different trial-types as in (C). **(J)** Average fluorescence traces in contra-coding ensembles on TS trials split by perceptual choice (top) and spatial choice (bottom). **(K)** Average choice probability (CP; top) and action probability (AP; bottom) scores in contra (red), ipsi (blue) and non-coding (grey) ensembles across the trial. Data show scores averaged across 52 sessions. Data in Figure 3 are recordings from 52 sessions across 13 mice (30 sessions in 7 symmetric-trained mice; 4.3 ± 2.0 (mean ± std) sessions per mouse, and 22 sessions in 6 asymmetric-trained mice; 3.7 ± 1.8 (mean ± std) sessions per mouse). Neural ensemble data were first averaged within session and then presented as the mean ± SEM across sessions.

We then imaged L2/3 during task performance (Figure 3C; n = 52 sessions in 13 mice; FOV depth: 150 – 200 µm; 256.5 ± 44.1 neurons per FOV; 109.9 ± 25.1 trials per experiment) and acquired videography to quantify task-evoked movement and licking (Figure S6). The 2P imaging field-of-view (FOV) was targeted to the stimulus ‘paddle’ responsive region of barrel cortex based on widefield fluorescence maps obtained at the start of each experiment (Figure S1C & S1D). As we did not find clear differences in neural responses across symmetric (n = 30 sessions in 7 mice; 4.3 ± 2 sessions per mouse; mean ± std) and asymmetric-trained mice (n = 22 sessions in 6 mice; 3.7 ± 1.8 sessions per mouse; Figure S7), we pooled both datasets unless otherwise stated.

### Sparse coding of contralateral and ipsilateral whisker information in L2/3

To isolate sensory-evoked signals prior to licking and reward, we analysed stimulus-evoked fluorescence (ΔF/F) responses during the first half of the delay epoch (Figure 3B *green shading;* 500 – 1000 ms post-stimulus). We confirmed this analysis window was free from licking using videography analysis (average video-detected RT post-stimulus = 1662.4 ± 142 ms; mean ± std; n = 52 sessions; Methods; Figure S6A & S6B). We first assessed whether neurons showed a preference for contra vs ipsi unilateral input by using Receiver Operating Characteristic (ROC) analysis to compute stimulus selectivity (Area Under the Curve; AUC). This revealed a broad distribution of selectivity across the L2/3 population, with some neurons preferring contra trials (red neurons in Figure 3D; neurons 1 and 2 in Figure 3E), and others preferring ipsi trials (blue neurons in Figure 3D; neurons 3 and 4 in Figure 3E). To identify statistically significant selective neurons, we compared contra and ipsi responses with a Wilcoxon rank-sum test (P < 0.05 = significant neuron). Surprisingly, a comparable proportion of L2/3 neurons were significantly selective for contra (fraction of all cells: 0.14 ± 0.05 (std); mean selectivity AUC 0.72 ± 0.04) and ipsi (fraction of all cells: 0.14 ± 0.07; mean selectivity AUC 0.31 ± 0.04) whisker input (Figure 3F & S7A). We refer to neurons with significant contra vs ipsi selectivity as ‘stimulus-coding’ and groups of stimulus-coding neurons in the FOV as ‘ensembles’. For the majority of our analyses, we averaged activity across neurons within ensembles, before comparing ensemble responses across sessions (average contra-coding ensemble size = 36.7 ± 15.8 neurons; ipsi-coding ensemble size = 37 ± 18 neurons; mean ± std; n = 52 sessions; 13 mice).

Increases in stimulus information in stimulus-coding ensembles were time-locked to stimulus onset (Figure 3G & S8A) and stimulus selectivity scores were similar irrespective of trial outcome (Figure S8B). Outside of the stimulus presentation window, quantification of whisking did not predict contra vs ipsi whisker stimulation (Figure S8D *bottom*), suggesting that active orofacial movements are similar across trial-types. Despite equal proportions of contra and ipsi-coding neurons in the FOV, unilateral responses were larger in contra-coding ensembles (preferred stimulus response (ΔF/F): contra-coding ensembles = 0.18 ± 0.11; ipsi-coding ensembles = 0.12 ± 0.06; ** P = 0.003; Wilcoxon signed-rank test; 52 sessions), consistent with an intrahemispheric bias for contralateral input. Across bilateral stimulation trials, responses in stimulus-coding ensembles increased as a function of preferred-stimulus intensity (Figure 3G & S7C). Stimulus-coding ensembles showed a small but significant decrease in fluorescence following presentation of the non-preferred stimulus (non-preferred stimulus response (ΔF/F): contra-coding neurons = −0.01 ± 0.03; ** P = 0.002; ipsi-coding neurons = −0.03 ± 0.03; *** P = 2.9 × 10^−8^; Wilcoxon signed-rank test difference tested from 0; n = 52 sessions, 13 mice). Most neurons did not have a statistically significant stimulus preference (fraction of all cells: 0.72 ± 0.09; 183 ± 37.9 cells per FOV; mean AUC 0.5 ± 0.02; Figure 3H & S7B) and did not show reliable responses on average across the task stimulus set (Figure 3I). We refer to these neurons as ‘non-coding’, however it is possible that these neurons code for other variables not explored under our task conditions. Our results therefore indicate that only a sparse subset of L2/3 neurons provide reliable stimulus-coding during task performance.

### L2/3 neurons do not predict categorical choice

During imaging experiments, we also included threshold stimulus (TS) trials, allowing us to correlate neural responses with behavioral decisions under stimulus uncertainty. As contra-coding ensembles showed stronger responses on bilateral trials, we first assessed if contra-coding ensembles predicted choice. Average fluorescence traces were similar on TS trials where mice reported different perceptual choices (Figure 3J *top*), and different spatial choices (Figure 3J *bottom*). To quantify this, we calculated choice probability^22^ (CP). Consistent with the stimulus-reward contingency, CP scores > 0.5 indicate increased fluorescence on trials with contra lickport choices for symmetric-trained mice, but ipsi lickport choices for asymmetric-trained mice. CP within stimulus-coding and non-coding ensembles remained at chance across the trial (mean delay-epoch CP in contra-coding ensembles = 0.5 ± 0.03; n.s. P = 0.51; ipsi-coding ensembles = 0.5 ± 0.03; n.s. P = 0.42; non-coding ensembles = 0.5 ± 0.01; n.s. P = 0.94; Wilcoxon signed-rank test difference tested from 0.5; Figure 3K *top* & S8E) with no clear correlation between stimulus-selectivity and CP scores across the L2/3 population (Figure S8C *top*). We also did not find obvious choice-coding in either symmetric or asymmetric-trained cohorts of mice (Figure S7A & S7D) and did not detect statistically significant choice neurons above the expected false positive rate (5%; fraction of neurons significant for contra perceptual choice: 0.05 ± 0.02; n.s. P = 0.92; ipsi perceptual choice: 0.05 ± 0.02; n.s. P = 0.61; Wilcoxon signed-rank test; difference tested against 0.05; Figure S8E). We then calculated a complementary metric we refer to as ‘action’ probability (AP), with AP scores > 0.5 predicting contralateral lickport choices and vice versa. Average AP scores also remained at chance (mean delay-epoch AP in contra-coding ensembles = 0.51 ± 0.07; n.s. P = 0.35; ipsi-coding ensembles = 0.5 ± 0.07; n.s. P = 0.8; non-coding ensembles = 0.5 ± 0.06; n.s. P = 0.9; Figure 3K *bottom* & S8C *bottom*). We also did not detect significant spatial choice-predictive neurons above chance (fraction of neurons significant for contra lickport choice: 0.06 ± 0.02; n.s. P = 0.3; ipsi lickport choice: 0.05 ± 0.02; n.s. P = 0.93; Figure S8F).

However, a small proportion of neurons predicted if mice would report a decision (‘lick’) vs miss (‘no lick’) in the trial (fraction of neurons significant for ‘lick’ trials: 0.07 ± 0.04; *** P = 1.5 × 10^−4^; Wilcoxon signed-rank test tested against 0.05; Figure S8G *left*). We summarised ‘lick’ vs ‘no lick’ discrimination by calculating detect probability (DP). DP was significantly above chance in stimulus-coding ensembles (DP: contra-coding ensembles; 0.53 ± 0.11; ** P = 0.002; ipsi-coding ensembles 0.52 ± 0.1; ** P = 0.009) but not in non-coding ensembles (DP in non-coding ensembles; 0.5 ± 0.09; n.s. P = 0.32). DP scores in stimulus-coding ensembles were larger than in non-coding ensembles (ΔDP = 0.02 ± 0.06; ** P = 0.008). However, performing the same ROC analysis on videography-extracted movement traces revealed that ‘lick’ trials could also be predicted from increases in whisking (Contra whisking DP: 0.58 ± 0.02; *** P = 4 × 10^−6^; Ipsi whisking DP: 0.56 ± 0.01; *** P = 7.9 × 10^−4^; Wilcoxon signed-rank test difference tested against 0.5; Figure S8G *bottom*) and body movement (Body movement DP = 0.57 ± 0.15; *** P = 8.4 × 10^−4^). Thus, it is possible that neural DP signals in the stimulus-coding ensembles reflect differences in non-instructed movements, as increases in active whisking will generate afferent sensory input into barrel cortex. In comparison, whisking quantified from either the contra or ipsi side of the snout did not predict contralateral perceptual choice (Contra whisking CP: 0.47 ± 0.12; n.s. P = 0.17; Ipsi whisking CP: 0.48 ± 0.11; n.s. P = 0.16; Wilcoxon signed-rank test difference tested against 0.5; Figure S8E *bottom*), or contralateral lickport choice (Contra whisking AP: 0.52 ± 0.12; n.s. P = 0.23; Ipsi whisking AP: 0.52 ± 0.11; n.s. P = 0.19; Wilcoxon signed-rank test difference tested against 0.5). This suggests that perceptual decisions and motor planning are not overtly reflected in lateralized orofacial movements. However, lateralized whisking did predict spatial choice during the choice window, consistent with an increase in ipsilateral orofacial movement during execution of bouts of directional licking (Figure S8F *bottom*). Thus, while barrel cortex showed sparse coding of stimulus information, we did not find a consistent predictive relationship between neural responses, or task-evoked movements, and categorical choice.

### Targeted two-photon photostimulation of L2/3 neurons during behavior

Our imaging results indicate that barrel cortex encodes contra and ipsi information in sparse subsets of neurons. While our experiments in the non-delay task show that manipulating barrel cortex drives lateralized biases in psychophysical performance (Figure 2), the number, functional identity, and layer-specificity of neurons activated by one-photon optogenetic stimulation are poorly controlled. To bridge this gap, we designed an all-optical experiment^35,53,54^ to test how targeted stimulation of different L2/3 ensembles impacted perceptual report of the whisker TS (Figure 4A-C). Following behavioral imaging (Figure 3), we selected two groups of 30 pyramidal neurons in the FOV co-expressing GCaMP6s and soma-targeted C1V1 (st-C1V1, Figure S1D) for holographic photostimulation (PS; Figure 4D). We designed target groups with the intention that one would have a contra whisker selectivity bias, while the other an ipsi bias, by selecting the 30 PS-responsive neurons with the highest and lowest stimulus-selectivity scores respectively (Figure S9A-C). However, as neurons in the FOV with strong responses to both PS and sensory input were rare (Figure S9D & S9E), our strategy likely includes a large number of target neurons with weak stimulus selectivity. PS was performed using a spatial light modulator (SLM^61^; Figure 4A) together with a galvanometer-based spiral scanning strategy^53^ (Methods). PS-evoked responses were quantified using simultaneous 2P calcium imaging (quantified 500 – 1000 ms post-stimulus; green shading in Figure 4C). We paired PS with the whisker TS (TS+PS; Figure 4C), and randomized PS across contra and ipsi-biased target groups across trials. We also delivered TS, PS, catch and unilateral whisker trials in a randomized order during the experiment.

**Figure 4.**
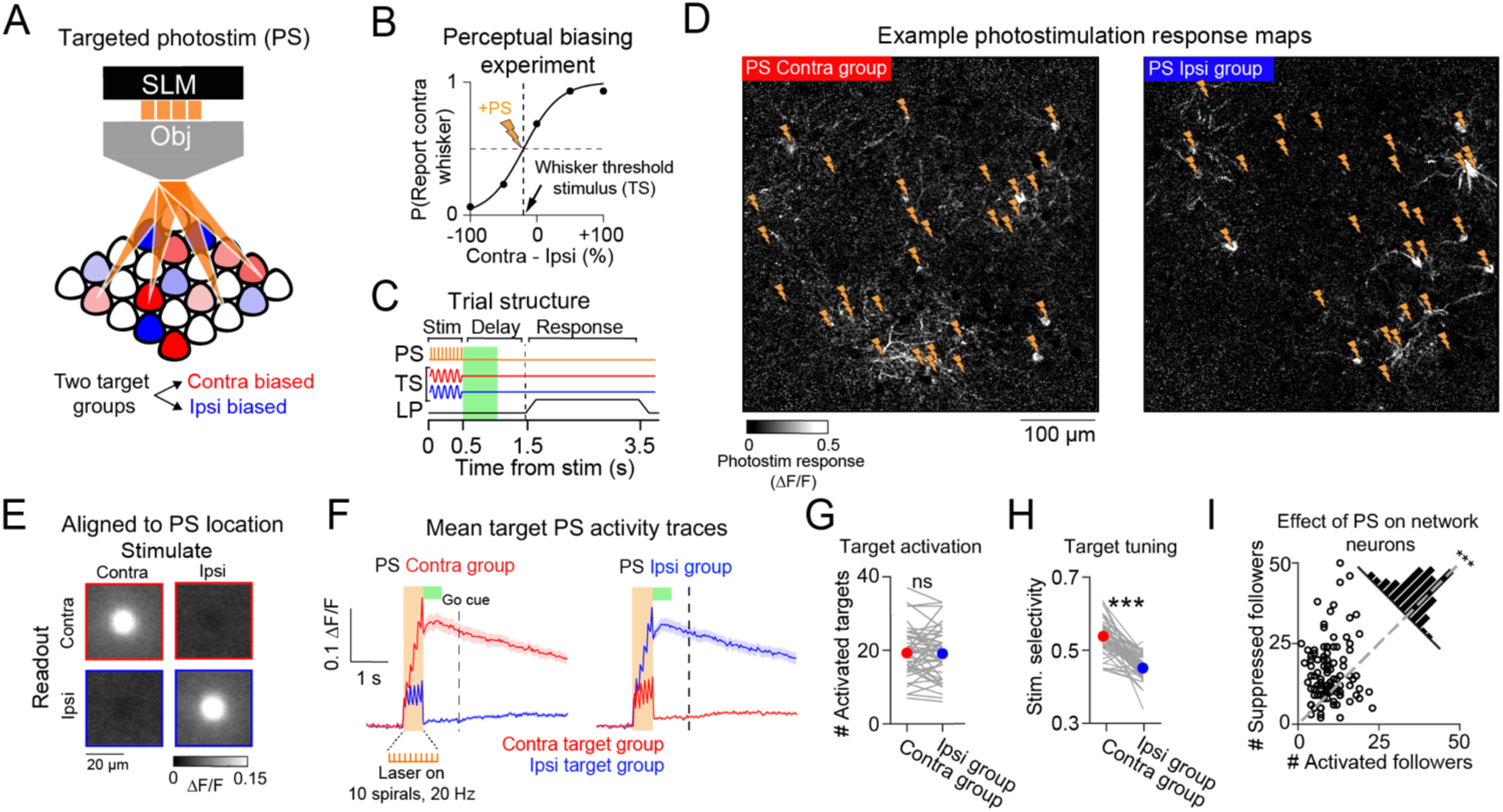
Two-photon photostimulation of L2/3 neurons during whisker discrimination. **(A)** SLM-targeted two-photon photostimulation (PS) of contra vs ipsi-biased L2/3 ensembles. **(B)** Photostimulation was delivered simultaneously with the whisker threshold stimulus (TS). **(C)** Schematic showing the trial structure for paired sensory and photostimulation (TS + PS) trials. The green shaded region shows the 2P imaging analysis window (500 - 1000 ms post-stimulus). **(D)** Photostimulation response maps showing the mean photostimulation-evoked response 500 - 1000 ms post-stimulus from an example session. Stimulation of the contra and ipsi target groups are shown on the left and right respectively, with lightning bolts indicating the targeted locations. **(E)** Average pixelwise PS response maps centred on all PS spiral sites for contra (top row) and ipsi (bottom row) SLM targets, on contra group stimulation trials (left column) and ipsi group stimulation trials (right column; average across 1560 target sites across 52 session). **(F)** Average fluorescence traces extracted from contra (red) and ipsi (blue) target neurons on photostimulation trials. The orange bar indicates the photostimulation duration, the green bar shows the response analysis window, and the vertical dashed line shows the go cue. Data are averaged across 52 sets of target groups from 52 sessions across 13 mice; 1992 activated target neurons in total. **(G)** Quantification of the number of target neurons activated by photostimulation during TS + PS trials. The coloured marker shows the mean and grey lines show individual session data. **(H)** Same as in (G) but showing quantification of the average stimulus-selectivity across activated target groups. **(I)** Quantification of the number of suppressed vs activated network followers during TS + PS trials. 52 sets of followers in 52 sessions; 13 mice; 1299 activated and 2078 suppressed follower neurons in total. Statistical comparisons were made across sessions with Wilcoxon signed-rank tests. * P < 0.05, ** P < 0.01; *** P < 0.001.

In post-hoc analysis, we parsed neurons into ‘target’ and ‘network’ categories based on lateral somatic proximity to the nearest PS spiral site (< 20 µm distance = ‘target’ neuron; Figure S10A-D). PS evoked soma-shaped increases in fluorescence at the intended spatial locations in the FOV (Figure 4D and 4E) and selective stimulation-locked fluorescence increases in the corresponding target groups (Figure 4F). We used a Wilcoxon rank-sum test (with a P < 0.05 criterion) to identify neurons with statistically significant PS-evoked changes in fluorescence and refer to neurons in target ‘zones’ with significant response increases as ‘activated’ targets. Despite delivering light to 30 target locations per stimulation pattern, PS activated a variable number of target neurons across different stimulation patterns (range 6 – 37 targets; activated targets in TS+PS contra group 19.3 ± 6.5; mean ± std; activated targets in TS+PS ipsi group = 19.1 ± 6.8; n.s. P = 0.72; Wilcoxon signed-rank test; n = 52 sessions; Figure 4G). Notably, the same photostimulus activated a larger number of targets when delivered in the absence of TS input (# activated targets: PS trials = 25 ± 6.3 neurons; TS+PS trials = 19.2 ± 6.6 neurons; *** P = 1.1 × 10^−15^; comparison across 104 target groups in 52 sessions). The number of activated targets on PS and TS+PS trials was highly correlated (Figure S10E) and was also correlated with the number of PS-responsive neurons in the FOV (Figure S10F), suggesting that variability in opsin expression across different experiments may influence the success rate of targeted photostimulation. Moreover, the number of activated targets was also predicted by variability in the total number of neurons located within target ‘zones’ across different stimulation patterns (Figure S10F), which we did not control for experimentally. Importantly, despite variability in target responses, activated target groups had a weak, but consistent, whisker stimulus selectivity bias (mean selectivity difference across groups = 0.083 ± 0.063, P = 5.3 × 10^−9^; Wilcoxon signed-rank test; Figure 4H). We also quantified the impact of targeted PS on non-targeted ‘network’ neurons. We refer to neurons in the local network that showed significant increases vs decreases in evoked fluorescence on TS+PS trials as activated vs suppressed ‘followers’, respectively. Across experiments we found that a larger number of followers were suppressed on TS+PS trials (# activated followers: 12.5 ± 5.6; # suppressed followers: 20 ± 11.5; *** P = 8.9 × 10^−8^; Wilcoxon signed-rank test; n = 104 photostimulation conditions; 52 sessions; Figure 4I). Thus, patterned 2P photostimulation resulted in selective activation of neurons within target zones while predominantly suppressing other neurons in the local L2/3 network.

### Perceptual bias scales with the number of photostimulated L2/3 neurons

Photostimulation did not result in average changes in perceptual choice tendency on TS trials (P(Report contra whisker) on TS trials: 0.56 ± 0.2; 45.7 ± 21.7 trials; mean ± std; on TS+PS contra trials: 0.56 ± 0.24; 22.8 ± 10.5 trials; n.s. P = 0.68; on TS + PS ipsi trials: 0.58 ± 0.24; 23.4 ± 10.8 trials; n.s. P = 0.33; Wilcoxon signed-rank test; n = 52 sessions; Figure 5A). Target group whisker selectivity also did not correlate with perceptual bias across sessions (ΔP(Report contra whisker) vs Target group selectivity; Pearson’s corr (r) = 0.05; n.s. P = 0.06; Figure 5B). Instead, we found that perceptual bias significantly correlated with the number of activated target neurons (ΔP(Report contra whisker) vs # activated targets; Pearson’s corr (r) = 0.3; ** P = 0.002; Figure 5C). This correlation was present when using different statistical thresholds for defining activated targets (Figure S11A) and was absent when we repeated our analysis on resampled control TS trials (average Pearson’s corr (r) on shuffled trials = 0.01 ± 0.1; mean ± std; 10000 shuffles; ** P < 0.002; Figure S11B). Moreover, target groups did not show choice probability on control trials that differed from chance (mean CP in contra-biased target group 0.5 ± 0.03; n.s. P = 0.67; ipsi-biased target group 0.51 ± 0.04; n.s. P = 0.56; Wilcoxon signed-rank test; Figure S11C), and target group CP did not predict the impact of photostimulation on behavior or the number of activated targets across sessions (Figure S11D).

**Figure 5.**
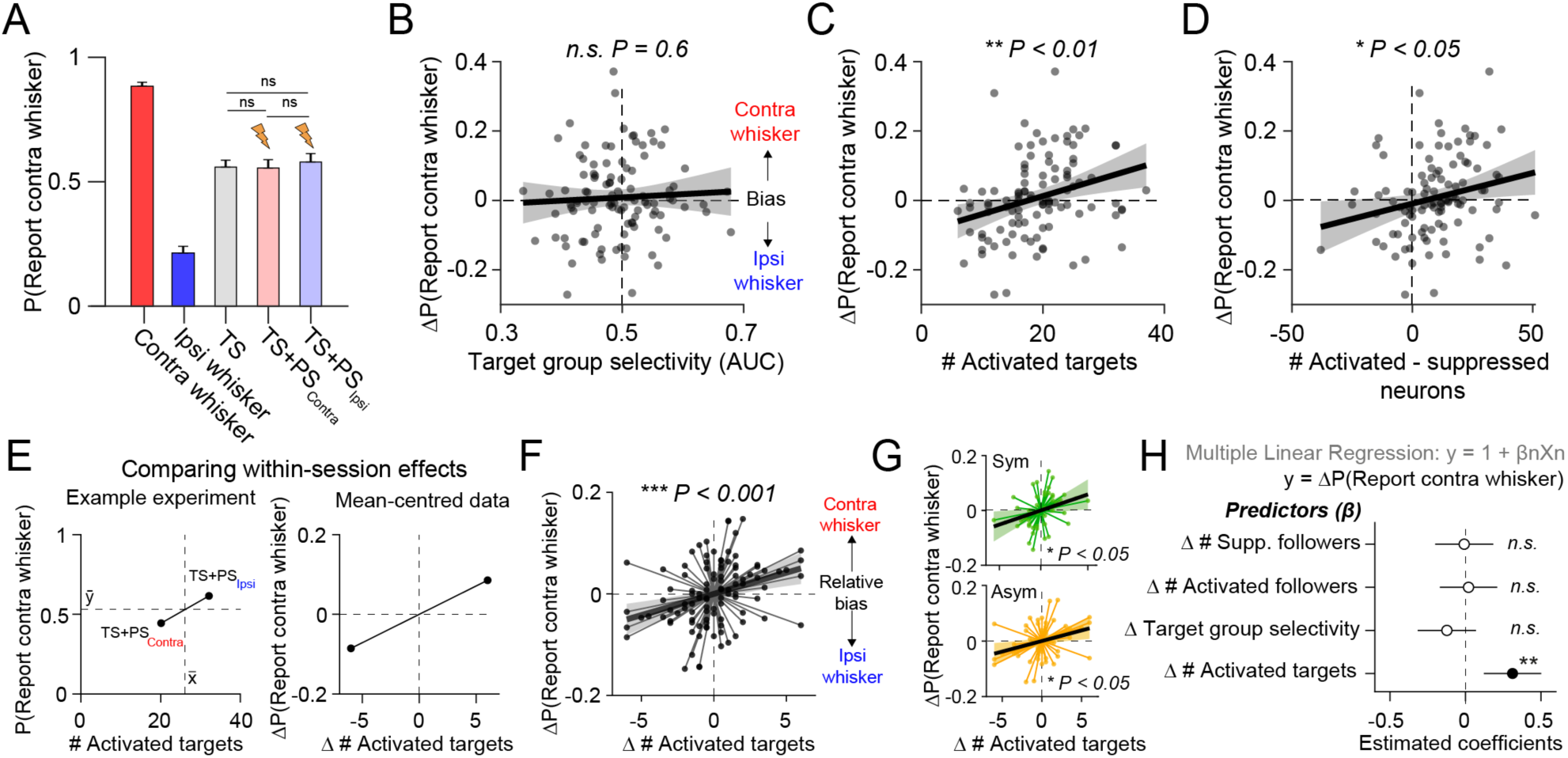
The number of activated target neurons predicts perceptual bias. **(A)** Quantification of average perceptual choice tendency across trial-types during the targeted photostimulation experiment. Photostimulation did not result in significant changes in discrimination across experimental sessions; Wilcoxon signed-rank tests. n.s. = not significant P > 0.05. **(B)** Comparison of perceptual bias and target group whisker selectivity (Pearson’s corr (r) = 0.05, n.s. P = 0.6). **(C)** Comparison of perceptual bias and the number of activated target neurons (Pearson’s corr (r) = 0.3, ** P = 0.002). **(D)** Comparison of perceptual bias and the net change in population activity (Pearson’s corr (r) = 0.23, * P = 0.02). **(E)** Within-session differences across photostimulation conditions were compared by ‘mean-centering’ the data-points from each session. This procedure is shown with a single example session. The dashed lines on the left plot indicate the mean P(Report contra whisker) (horizontal) and target response (vertical) across the two conditions. **(F)** Correlation between within-session difference in the number of activated targets and perceptual bias across all mean-centred data points (Pearson’s corr (r) = 0.33; *** P = 0.0007). Data points corresponding to the same session are joined with a thin line that passes through the origin. **(G)** The same as in (F) but split by sessions from symmetric (green; top; Pearson’s corr (r) = 0.33; * P = 0.011; n = 30 exp) and asymmetric (orange; bottom; Pearson’s corr (r) = 0.34; P = 0.025; n = 22 exp) trained mice. **(H)** A multiple linear regression (MLR) model was used to summarise the relationship between target and network photostimulation predictors and within-session perceptual bias. Statistically significant predictors are shown with black markers and non-significant predictors with white markers, with error bars indicating 95% coefficient confidence intervals. Data in Figure 5 are from 52 sessions across 13 mice. Each data point represents an individual photostimulation condition (one for TS + PS contra trials and one for TS + PS ipsi trials), with a total of 104 data points. Total number of control TS trials = 2374, total number of TS trials with PS = 2403. In correlation plots, shaded error bars denote the 95% confidence bounds of a linear regression fit to the data (black line).

Intriguingly, activating low numbers of targets appeared to bias choice towards the ipsilateral whisker (Figure 5C). By quantifying the number of activated vs suppressed neurons across both targets and followers, we found that perceptual bias also correlated with net change in population activity (ΔP(Report contra whisker) vs # activated – suppressed neurons; Pearson’s corr (r) = 0.23; * P = 0.02; Figure 5D). This suggests that ipsilateral biases tended to occur in sessions where photostimulation was comparatively weak, and thus the net effect on the L2/3 circuit was slightly suppressive (Figure 5D). However, quantification of follower neurons alone was not sufficient to predict changes in behavior (Figure S11E), implying that the impact of PS on perceptual bias cannot be wholly explained by differences in network state.

### Within-session perceptual bias correlates with the number of activated target neurons

As we stimulated two target groups in each session, we then analysed whether differences in the number of activated target neurons across target groups explained variability in within-session perceptual bias. To examine this effect, we mean-centred the data-points from the two photostimulation conditions from each session (Figure 5E). Across the mean-centred data set, variability in the number of activated targets also significantly correlated with relative perceptual bias (ΔP(Report contra whisker) vs Δ # activated targets; Pearson’s corr (r) = 0.33; *** P = 0.0007; Figure 5F). This correlation was present separately in symmetric (Pearson’s corr (r) = 0.33; * P = 0.011; n = 30 sessions, 7 mice; Figure 5G *Top*) and asymmetric-trained cohorts of mice (Pearson’s corr (r) = 0.34; * P = 0.025; n = 22 sessions, 6 mice; Figure 5G *bottom*). We then used a multiple linear regression model to summarise within-session effects of photostimulation on perceptual choice bias (Figure 5H). Perceptual bias was significantly predicted by the number of activated targets (estimated coefficient = 0.31; *t*-stat = 3.3; ** P = 0.001) but not the number of activated (estimated coefficient = 0.03; *t*-stat = 0.32; n.s. P = 0.74) or suppressed (estimated coefficient = −0.01; *t*-stat = −0.5; n.s. P = 0.96) follower neurons. Within-session perceptual bias was also not correlated with variability in target group whisker stimulus selectivity (estimated coefficient = −0.16; *t*-stat = −1.6; n.s. P = 0.1), further indicating that the perceptual effect of L2/3 photostimulation is predicted by the number, and not tuning, of activated target neurons. Additional analysis confirmed that neither within-session perceptual bias nor the number of activated target neurons, correlated with differences in pre-stimulus network fluorescence or peri-stimulus variability in whisking or body movement across photostimulation trial-types (Figure S11F). Thus, our results indicate that sparse unilateral manipulation of the L2/3 circuit during bilateral discrimination causally drives a lateralized perceptual bias that scales with the number of neurons activated.

### Activation of task-silent target neurons predicts perceptual bias

In contrast with previous studies^41,44,45,62^, we did not find behavioral changes consistent with the tuning of the stimulated neurons (Figure 5B & 5H). However, while the target groups did show an average stimulus-selectivity bias (Figure 4H), additional analysis revealed that the majority of activated targets did not have a statistically significant whisker stimulus preference (Figure 6A). As stimulation predominantly activated non-coding target neurons in the FOV, we assessed whether this was sufficient to predict the perceptual effect of targeted photostimulation. A multiple linear regression analysis (Figure 6B) confirmed that the number of non-coding target neurons activated significantly predicted perceptual bias (estimated coefficient = 0.3; *t*-stat = 3.14; ** P = 0.002). In comparison, neither the number of activated contra-coding targets (estimated coefficient = 0.05; *t*-stat = 0.46; n.s. P = 0.64) nor the number of ipsi-coding targets (estimated coefficient = 0.03; *t*-stat = 0.32; n.s. P = 0.75) predicted the behavioral effect. The correlation between non-coding target neuron activation and perceptual bias was present across sessions (Pearson’s corr (r) = 0.31; ** P = 0.001; Figure S12A) and within-session (Pearson’s corr (r) = 0.21; * P = 0.03; Figure S12B).

**Figure 6.**
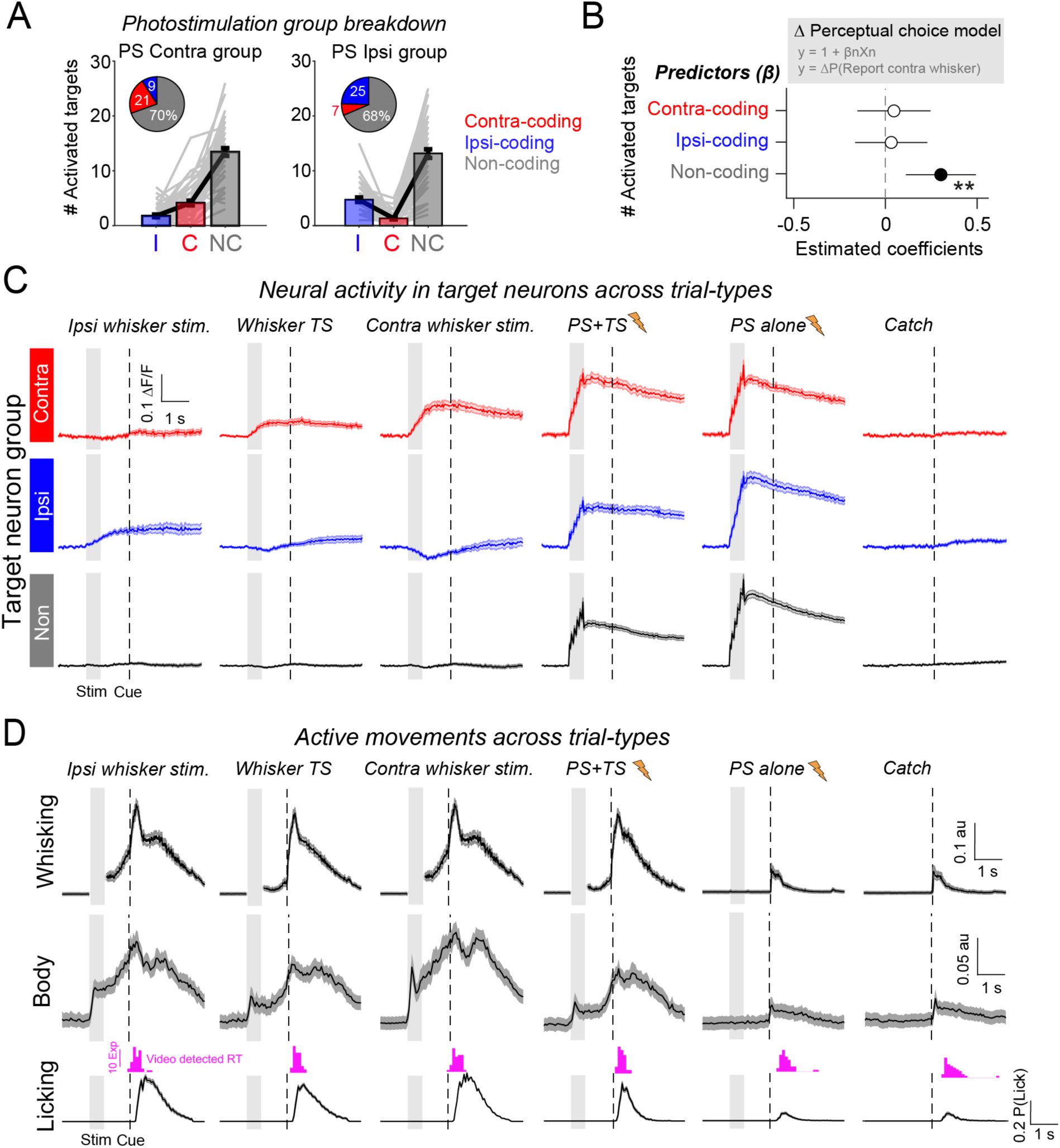
Photostimulation of non-coding neurons predicts perceptual bias. **(A)** Quantification of the number of contra-coding (red), ipsi-coding (blue) and whisker non-coding (grey) activated target neurons that made up the two photostimulation target groups. Individual lines show individual sessions. The inset circle charts indicate the average proportional group summary. **(B)** A multiple linear regression (MLR) model summarised the relationship between the number of photostimulated stimulus-coding and non-coding neurons and perceptual bias. Marker points shown the estimated coefficients with error bars indicating the 95% confidence intervals. **(C)** Trial-evoked activity traces from contra-coding (red), ipsi-coding (blue) and non-coding (grey) photostimulation target neurons are shown across different trial-types. Data show the average target group activity across sessions. The grey shaded bar shows the duration of the stimulus, and the vertical dashed line shows the time of the go cue. **(D)** The time course of average trial-evoked contralateral whisking (top), body movement (middle) and licking (bottom) behavioural measures are shown across trial-types. Whisking and body movement were assessed with videography, licking traces were assessed using the electrical lickport detector. The magenta histogram shows the distribution of mean reaction times assessed using videography. Note that during whisker stimulation (grey shading), we do not plot whisking traces as it is unclear which whisker movements are driven by the stimulus and which are the result of active movement. Data in Figure 6 are from 52 sessions in 13 mice. Total number of neurons analysed: 274 contra-coding targets; 329 ipsi-coding targets; 1381 non-coding targets. Data are shown as the mean and SEM across sessions.

Non-coding target neurons showed reliable photostimulation-evoked responses but did not show whisker-evoked responses (Figure 6C). This was consistent for non-coding target neurons that were included in the contra-biased and the ipsi-biased target groups (Figure S13). Analysis of behavioral videography also revealed that trials with whisker stimulation evoked sustained increases in active whisking, body movement and licking (Figure 6D & S6C). As fluorescence in non-coding target neurons remained at pre-stimulus levels throughout the trial-epoch, this indicates that the non-coding target neurons are also not reliably activated by instructed or non-instructed movements during task performance. Our findings therefore indicate that small numbers of ‘task-silent’ L2/3 neurons, which show no clear functional relationship to sensory, motor or reward variables during task performance, can influence perceptual decisions if recruited into the active population through targeted optogenetic manipulation.

### Patterned stimulation reveals specificity of L2/3 inhibition during whisker processing

Comparing target responses across trial-types revealed that photostimulation responses were notably larger in the absence of concurrent whisker stimulation, particularly in ipsi-coding and non-coding target neurons (Figure 6C). This prompted us to examine the interaction between sensory and photostimulation triggered activity patterns across the FOV (Figure 7A). For some targets in the FOV, PS responses were near abolished on TS+PS trials, and all photostimulation target site locations showed a reduction in fluorescence when comparing PS trials with TS+PS trials (Figure 7B). To assess the relationship between the decrease in PS response and the TS-evoked sensory response, we quantified the average difference in the target group response on PS and TS+PS trials and compared this with the average response to TS input in the TS-responsive non-targeted ‘network’ neurons (Figure 7C). Across sessions, the change in target group PS response was negatively correlated with the network response to sensory input (Pearson’s corr (r) = −0.49; *** P = 1.3 × 10^−7^; Figure 7D). This indicates that the stronger the representation of whisker input in the L2/3 network, the stronger the suppressive effect on photostimulation-evoked activity in the target neurons.

**Figure 7.**
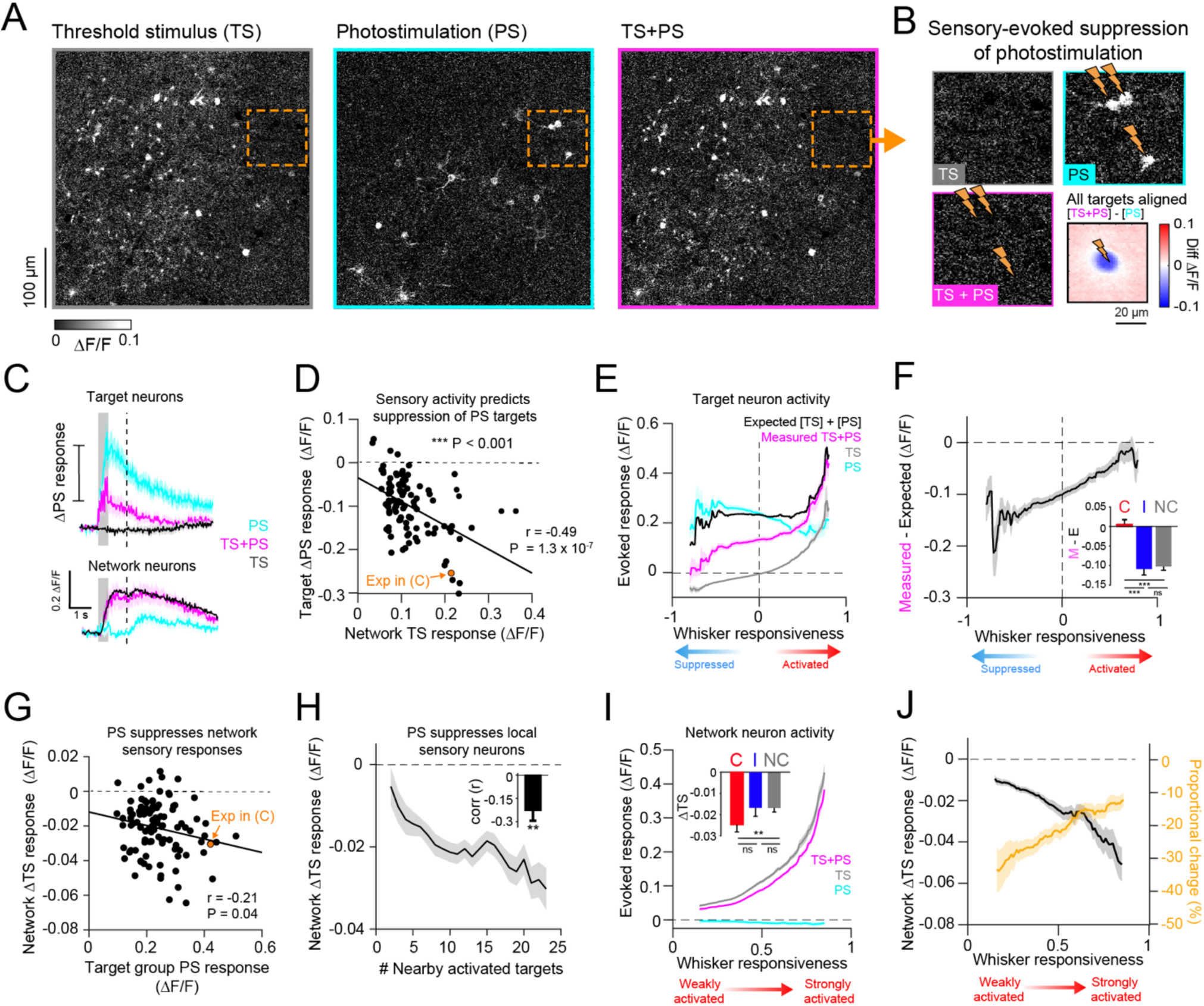
Patterned photostimulation reveals potent inhibitory pressure in L2/3 during task performance. **(A)** Response maps for threshold stimulus (TS; grey), photostimulation (PS; cyan) and combined TS and PS (magenta) trials for an example sessions. Responses were averaged 500 - 1000 ms post-stimulus. **(B)** Close up of the orange-bounded region shown in (A). Lightning bolts indicate the location of PS targets. Bottom right shows the average pixel-wise fluorescence difference across TS+PS and PS trials centre-aligned on all photostimulation target locations (average across 104 photostimulation target groups in 52 sessions). **(C)** Extracted fluorescence traces from target neurons (top; n = 23 neurons) and TS responsive network neurons (bottom; n = 57 neurons) across trial-types from the example sessions shown in (A). **(D)** Suppression of photostimulation responses in the target neuron group on PS vs TS+PS trials is correlated with the mean response to TS whisker trials in non-targeted network cell group (Pearson’s corr (r) = −0.49; *** P = 1.3 × 10^−7^). Each data point represents a single session. **(E)** Responses in photostimulation target neurons across different trial-types are plotted as a function of whisker responsiveness. **(F)** The difference between the measured (magenta in E) and expected (black in E) response to combined TS and PS stimuli in target neurons is plotted as function of whisker responsiveness. The inset shows quantification of this difference averaged across contra-coding (red), ipsi-coding (blue) and non-coding (grey) target neuron ensembles. **(G)** Suppression of TS responses in network neuron groups on TS+PS trials is plotted against average PS-evoked responses in target neuron groups (Pearson’s corr (r) = −0.21; * P = 0.04). **(H)** Average PS evoked suppression of TS responses in network neurons is binned as a function of the number of nearby activated target neurons (counted within a 200 µm radius). The inset shows the average Pearson’s correlation coefficient between sensory suppression and number of nearby targets across sessions (r = −0.21 ± 0.47; mean ± std; ** P = 0.002 Wilcoxon signed rank test tested against 0). **(I)** Responses in TS-responsive network neurons across trial-types are plotted as a function of whisker-responsiveness. The inset shows quantification of the change across TS and TS+PS trials with respect to ensemble groups as in (F). **(J)** The difference in network neuron response on TS and TS+PS is shown as a function of whisker responsiveness. The absolute change is shown in black, and the proportional change relative to TS baseline is shown in orange. Data in Figure 7 come from 104 photostimulation conditions across 52 sessions in 13 mice. Total number of target neurons analysed = 1992, total number of network neurons analysed = 2429. Total number of trials analysed: 3466 TS, 3568 PS; 3447 TS+PS trials. Data are shown as the mean with error bars showing SEM across sessions. Statistical comparisons were Wilcoxon signed-rank test. n.s. P > 0.05; * P < 0.05; ** P < 0.01; *** P < 0.001.

To examine whether this effect showed specificity at the single neuron level, we quantified fluorescence responses across trial-types in all activated targets as a function of whisker responsiveness (Figure 7E). We defined whisker responsiveness as the AUC score from an ROC analysis comparing TS responses with catch trial responses, and adjusted scores between −1 and 1 such that the sign denoted negative vs positive modulation by TS input. We then calculated the difference between measured TS+PS responses and the expected linear sum of separate TS and PS trial-evoked responses as an approximation for sensory-induced suppression of photostimulation activity (Figure 7F). Target neurons with high whisker responsiveness scores appeared largely unaffected by sensory-induced network suppression. In contrast, neurons with lower, or negative, whisker responsiveness scores showed a large difference in photostimulation response. When quantifying this effect with respect to stimulus-coding groups, we found that contra-coding targets groups showed no difference between measured and expected response (0.004 ± 0.01; n.s. P = 0.78), whereas both ipsi-coding target groups (−0.11 ± 0.11; *** P = 2.6 × 10^−8^) and non-coding target groups (−0.1 ± 0.07; *** P = 6.3 × 10^−10^) showed a strong suppression of photostimulation activity on TS trials (Figure 7F *inset*). Thus, patterned photostimulation revealed recruitment of potent cortical inhibition in L2/3 during whisker processing that appeared to preferentially impact ipsi-coding and non-coding neurons in the L2/3 circuit.

### Patterned stimulation suppresses local whisker-evoked signals

We next investigated the impact of patterned photostimulation on whisker-evoked responses in the local circuit. We focused our analyses on TS-responsive neurons that were not targeted for photostimulation. On average, targeted photostimulation reduced the amplitude of the TS-response in network neuron groups (ΔTS response = −0.02 ± 0.015 ΔF/F; *** P = 9.7 × 10^−18^, −19.4 ± 13.6% proportional change; n = 104 photostimulation conditions, 52 sessions). This reduction was correlated with the amplitude of photostimulation response in the target groups (Pearson’s corr (r) = −0.21; * P = 0.04; Figure 7G). The magnitude of sensory response suppression was stronger for network neurons in close proximity to large numbers of PS-activated target neurons (Figure 7H), suggesting that the suppressive effect of PS on the local L2/3 circuit reduces with increasing inter-somatic distance. All non-targeted TS-responsive neurons showed reduced TS responses on TS+PS trials on average (Figure 7I), with those that were part of contra-coding ensembles tending to show larger absolute changes in ΔF/F (Figure 7I *inset*). However, neurons with larger whisker responsiveness scores showed smaller proportional changes relative to the baseline amplitude of the TS-evoked response (Figure 7J). As strongly whisker-responsive neurons were relatively less inhibited by network-evoked suppression, this indicates that global inhibition might serve an important role in selectively eliminating weak responses in the circuit.

### Competitive interactions between contralateral and ipsilateral whisker signals in L2/3

Finally, to investigate the relationship between cortical inhibition and bilateral sensory coding, we performed additional experiments to examine interactions between contra and ipsi-evoked signals in L2/3. After performing targeted photostimulation, we mapped passive neuronal responses to a ‘3 × 3’ stimulus set of unilateral and bilateral stimuli (Figure 8A & 8B). Contra-coding ensembles, defined based on activity during the behavioral experiment, showed large responses to unilateral contralateral stimulation that appeared robust to the presence of concurrent ipsilateral whisker stimulation (Figure 8B *top*). In contrast, ipsilateral-evoked signals in ipsi-coding ensembles were weaker and were strongly suppressed by contralateral whisker input (Figure 8B *bottom*). To examine this relationship across the L2/3 population, we compared responses to the preferred unilateral stimulus (100% intensity) to bilateral stimulation (100% intensity on both sides) as a function of stimulus selectivity (Figure 8C). We then calculated the difference between the bilateral and preferred unilateral response (Figure 8D). Contra-coding ensembles showed a modest reduction in response on bilateral stimulation trials (change in response (ΔF/F): −0.01 ± 0.02; ** P = 0.001; Wilcoxon signed-rank test; n = 52 sessions), whereas ipsi-coding ensembles showed more dramatic attenuation (change in response (ΔF/F): −0.04 ± 0.03; *** P = 3.7 × 10^−9^; Figure 8D *inset*). The difference between contra and ipsi-coding ensemble suppression was significant (*** P = 1.9 × 10^−5^).

**Figure 8.**
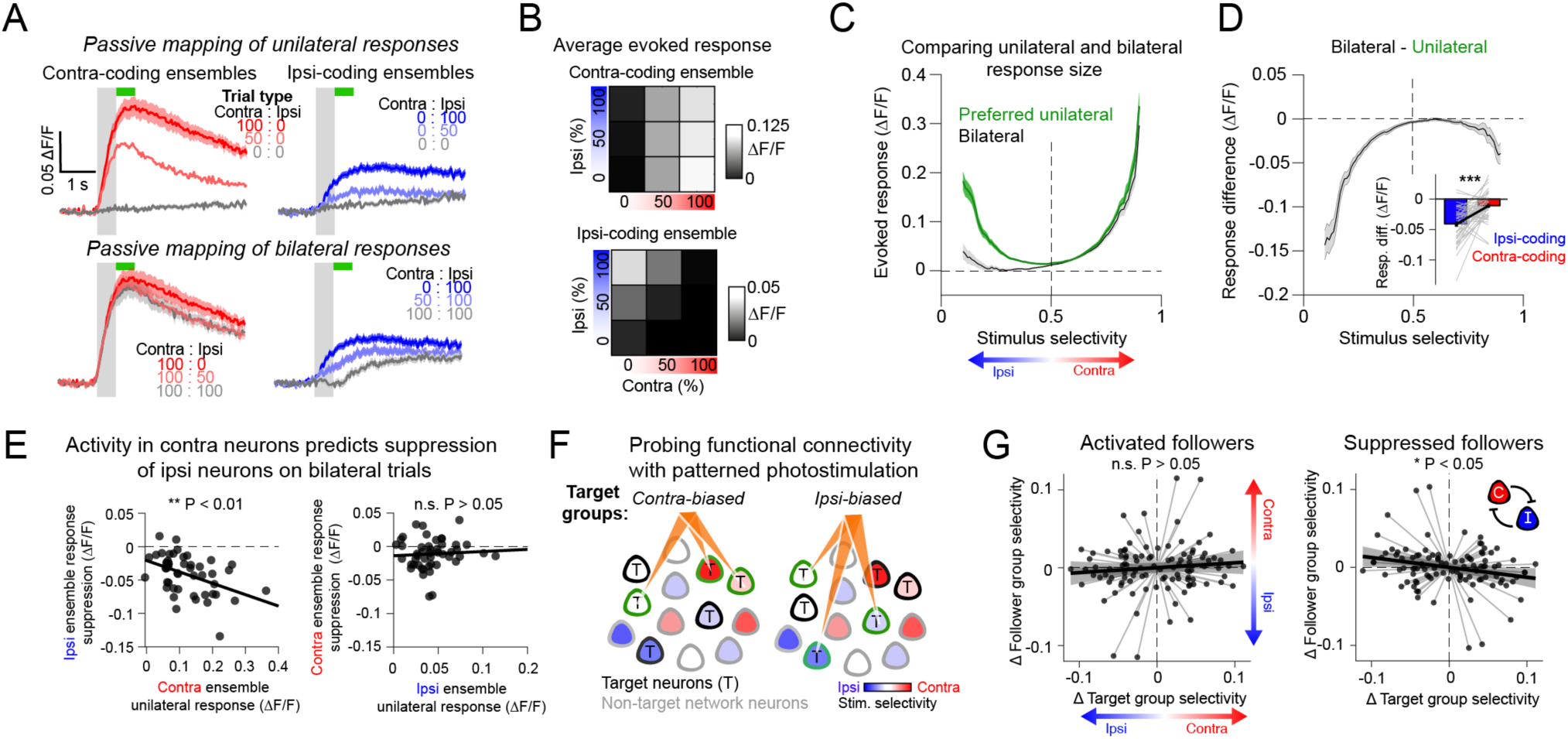
Antagonistic interactions between contra and ipsi-coding neurons in L2/3. **(A)** Evoked fluorescence responses in contra-coding (left; red) and ipsi-coding (right; blue) ensembles across a range of unilateral (top) and bilateral (bottom) whisker deflection intensities (%) during passive imaging sessions. **(B)** Quantification of the mean fluorescence responses across a ‘3 × 3’ matrix stimulus set in contra-coding (top) and ipsi-coding (bottom) ensembles (average ensemble response across 52 sessions). **(C)** Comparison of neuronal responses to preferred unilateral whisker stimulation (green; 100% unilateral stimulation) and matched bilateral whisker stimulation (black; 100% bilateral stimulation) plotted as a function of stimulus selectivity (average across 52 session). **(D)** The difference in unilateral and bilateral responses (shown in C) is plotted as a function of stimulus selectivity. The inset shows quantification of this difference in contra and ipsi-coding ensembles (n = 52 sessions; Wilcoxon signed-rank test; *** P < 0.001; thin grey lines show individual sessions). **(E)** Left: Across sessions, bilateral suppression of the ipsi-coding ensemble is correlated with the contra-coding ensemble response to contra whisker stimulation (Pearson’s corr (r) = −0.44; ** P = 0.002; n = 52 sessions). Right: Suppression of contra-coding ensemble did not correlate with ipsi ensemble activity (Pearson’s corr (r) = 0.07; n.s. P = 0.66). Each marker point represents an individual sessions, 52 sessions in total. Total number of contra-coding neurons analysed 1907; ipsi-coding neurons 1925. **(F)** Probing functional connectivity in the circuit using targeted photostimulation of whisker-biased target groups. **(G)** Comparing average whisker selectivity of network followers in response to targeted photostimulation. Left: Difference in target group whisker selectivity across photostimulation conditions does not correlate with average difference in selectivity of activated network followers (Pearson’s corr (r) = 0.1; n.s. P = 0.32). Right: Differences in target group whisker selectivity negatively correlates with whisker selectivity of negative followers (Pearson’s corr (r) = −0.23; * P = 0.018). Correlation is based on 104 mean-centred data points (2 photostimulation conditions per session, 52 sessions in total). Shaded error bars denote the 95% confidence bounds of a linear regression fit to the data (black line).

Across sessions, the bilateral suppression of the ipsilateral ensemble was correlated with the contra-coding ensemble response to unilateral contralateral stimulation (Pearson’s corr (r) −0.44; ** P = 0.002; Figure 8E *left*). The inverse relationship between contra-coding ensemble suppression and the ipsi-coding ensemble response was not significant (Pearson’s corr (r) 0.07; n.s. P = 0.66; Figure 8E *right*). This indicates that contra-evoked signals may dominate during bilateral whisker processing by suppressing ipsi-responses in the L2/3 circuit. To directly examine this relationship, we performed additional analysis of our photostimulation dataset to assess whether the whisker-selectivity of the PS target groups correlated with the selectivity of recruited ‘follower’ neurons in the local circuit. For each session, we restricted our analysis to neurons in the local network that were not part of either activated target group, and thus we could directly compare the local circuit response to two different photostimuli (Figure 8F). Across the dataset, variability in target group whisker selectivity did not correlate with variability in the average whisker selectivity of the activated follower group (Pearson’s corr (r) = 0.1; n.s. P = 0.32; Figure 8G *left*). However, target group whisker selectivity negatively correlated with the selectivity of the suppressed follower group (Pearson’s corr (r) = −0.23; * P = 0.018; Figure 8G *right*), indicating that stimulation of L2/3 contra-coding neurons tends to suppress ipsi-coding neurons in the FOV and vice versa. Thus, our findings demonstrate that strong competitive interactions between different L2/3 ensembles dominate sparse cortical dynamics during sensory processing and provide evidence that mutual inhibition between contra and ipsi-coding ensembles is a prominent feature of intrahemispheric L2/3 activity during cross-hemispheric cortical processing.

## DISCUSSION

We used two-photon calcium imaging and two-photon holographic photostimulation to probe sparse coding in barrel cortex. Our experiments demonstrate that sparse coding of whisker stimuli is ensured by preferential sensory-evoked inhibitory suppression of non-coding neurons, which may help to ensure high signal-to-noise ratio for detection of small perturbations. Moreover, while contralateral and ipsilateral whisker signals were represented in sparse subsets of stimulus-coding L2/3 neurons during task performance, the effect of intrahemispheric photostimulation on lateralized perceptual bias depended on the number of non-coding target neurons activated. This suggests that non-coding neurons can be engaged by the circuit to enhance sensory processing, which could be achieved endogenously via cortical plasticity^20,49^.

### A whisker discrimination task to probe tactile perception

We developed a behavioral task that meets all the necessary criteria for quantifying the link between neural circuit activity in sensory cortex and behavior: precise control of the sensory stimulus, temporal separation of sensation and action epochs, interpretable readout of behavior, and compatibility with cellular-resolution recording and manipulation techniques^63^. Through both gain- and loss-of-function perturbations, we demonstrate that the perceived intensity of contralateral whisker input is coupled to intrahemispheric barrel cortex activity. By manipulating the bilateral stimulus interval in the non-delay task, we also found that unilateral percepts were formed rapidly and showed robustness to late-arriving distraction characteristic of choice-related attractor state dynamics^64^. The barrel cortex-dependent sensory integration window, which we estimate to be ~100 ms based on temporal discrimination and photoinhibition experiments, is comparable to cortical processing windows in other whisker tasks^24,65,66^ and tasks using other sensory modalities^67^. It is also consistent with findings showing that whisker input, and microstimulation-induced activity, quickly spreads from barrel cortex to high-order sensory and motor areas^68,69^, and demonstrates that barrel cortex plays a causal role in initiating rapid sensorimotor transformations during goal-directed behavior.

Imaging barrel cortex also revealed robust ipsilateral coding of whisker stimulation, a functional property of somatosensory cortex that has not been previously investigated in the context of a rigorous head-fixed perceptual task. However, despite coding of bilateral stimulus information, we did not find strong evidence of choice-encoding in barrel cortex. On the one hand, choice could be represented in membrane potential dynamics^24^ or temporal spike patterns^70^, which may be difficult to resolve using calcium imaging, or in deeper cortical layers, which we did not sample during our experiments. Alternatively, decision-related processing may occur downstream from S1, in areas that also integrate bilateral whisker input such as striatum^71^ and/or S2^72,73^. Future work could investigate the downstream circuits implicated in the decision process, as well as compare the distinct neural pathways engaged during symmetric and asymmetric whisker-guided responses, to further elucidate the role of barrel cortex in coordinating cross-hemispheric sensorimotor behavior.

### Inter-hemispheric processing in barrel cortex

Cross-hemispheric projections are an important component of many cortical circuits^74,75^, and L2/3 is a prominent locus for sending and receiving callosal projections^76,77^. Our findings reveal several interesting features of cross-hemispheric processing in barrel cortex. For instance, our results indicate that L2/3 contains equal proportions of ipsilateral and contralateral whisker-selective neurons, and that strong competitive interactions between bilateral tactile signals occur at the level of primary somatosensory cortex during perceptual decision-making. This extends previous work characterising bilateral tactile responses in barrel cortex, which has predominantly been performed under anaesthesia or in awake but non-behaving states^73,78,79^. While ipsilaterally-tuned neurons showed robust activity on unilateral trials, their responses were strongly attenuated by concurrent contralateral whisker stimulation. In contrast, contra-evoked signals showed milder suppression due to ipsilateral input. As such, L2/3 activity was markedly biased towards contralateral sensory input on bilateral trials. Our findings support previous evidence that the short cross-callosal latency makes ipsilateral signals susceptible to rapid feedforward recruitment of cortical inhibition by contralateral-evoked thalamocortical input during simultaneous bilateral stimulation^78–82^, further indicating that the integration of bilateral tactile signals may relate to the precise spatiotemporal sequence of whisker stimulation^78,79^.

In addition to antagonistic intra-cortical signals, our behavioral experiments also indicate that competition between hemispheres plays an important role in task-related perceptual processing. For example, results from the one-photon optogenetic-biasing experiments (in the non-delay task) show that manipulating one hemisphere has an equal but opposite effect on the reaction time for the non-stimulated hemisphere. The push-pull direction of the reaction time bias switches depending on whether one delivers unilateral photoinhibition or photoactivation, providing strong causal evidence that both hemispheres have the capacity to directly compete during task performance^83–86^. Moreover, mice trained on the non-delay task showed remarkable sensitivity to temporal offsets in bilateral whisker stimulation. This could also be the result of interhemispheric inhibition, as rapid suppression of the contralateral hemisphere by the leading whisker side would thus attenuate the late-arriving perceptual signal in the ipsilateral hemisphere.

In rodent cortex, several mechanisms have been identified that mediate inhibitory interactions between hemispheres. These include transcallosal feedforward recruitment of local PV interneurons across layers^73,77,86,87^, recruitment of interneurons in superficial layers, which subsequently inhibit the distal dendrites of neurons in deeper layers^85^, as well as direct suppression of pyramidal neurons via callosally-projecting inhibitory neurons^88^. This demonstrates that that the contralateral hemisphere can exert significant suppressive influence on intracortical sensory processing via a diverse range of mechanisms. However, the extent to which these different inhibitory pathways are engaged in parallel, or differentially under stimulus-specific or task-specific conditions, remains unclear. Moreover, callosal projections can also mediate more refined functions including the homotopic transfer of sensorimotor signals and learned information^74,89,90^, and shaping receptive fields and circuit plasticity^83,84,91^. Future work is therefore needed to rigorously examine how organization of callosal microcircuits shapes bilateral somatosensory perception, learning and memory.

### Intracortical inhibition enforces sparse coding in L2/3

Our experiments revealed strong antagonism between different populations of sensory-selective neurons, as well as competition between photostimulation and sensory-evoked signals in L2/3. This provides strong evidence that recruitment of inhibition is the basis of competitive interactions between different cortical excitatory ensembles during awake behaving states^14,15,92^, and demonstrates the utility of combining patterned optogenetic stimulation and population calcium imaging to probe functional dynamics in neural circuits^62,93,94^. Targeted photostimulation responses were strikingly attenuated when paired with simultaneous whisker deflection. This suggests that strong inhibitory mechanisms are engaged during task performance to balance network excitation, which under our task conditions could accumulate across feedforward^12,13,80,81,95^, feedback^96^, lateral^97,98^, and inter-hemispheric^85^ sources.

In line with other recent studies, we also found that targeted stimulation of pyramidal neurons suppressed other non-targeted pyramidal neurons in the local L2/3 circuit^35,93,98,99^. This provides further evidence that dense connectivity between excitatory and inhibitory neurons mediates strong lateral competition in cortex^8,82,98–100^ and that the recruitment of local inhibitory mechanisms scales with the number of concurrently activated pyramidal neurons^35,92,97^. Moreover, we present evidence that contra and ipsi-coding neurons exhibit signatures of intra-cortical mutual inhibition – suggesting that specific organisation of inhibitory microcircuits in L2/3 may support enhanced intrahemispheric discrimination of bilateral tactile space.

More generally, our analysis of the inhibitory interactions between photostimulation and sensory-evoked ensembles indicates that not only does strong inhibitory pressure play a major role in enforcing the sparsity of cortical responses^14,82,92^, but that sensitivity to inhibitory pressure appears non-uniformly distributed across the population. Photostimulation-evoked suppression of network neurons had the greatest proportional effects on responses that were initially small, consistent with previous findings indicating that disproportionate effects of cortical inhibition across heterogenous populations of barrel cortex neurons may serve to enhance sparse whisker coding^97,101^. This suggests that interactions between excitatory and inhibitory microcircuits exhibit a specific structure to enhance the signal-to-noise ratio of robust contralateral sensory signalling while maintaining sparsity of responses. Our findings also imply that weak sensory responses are more easily overridden by global inhibition than strong sensory responses, making strong responses stand out in a competitive environment to be picked up by the readout mechanism and become perceptually relevant. This is also consistent with demonstrations that a threshold level of excitation is required overcome strong inhibitory mechanisms in L2/3 to drive perceptually salient changes in cortical activity^35,41^.

### Small numbers of non-coding neurons can contribute to perception

We demonstrate that the targeted manipulation of surprisingly few L2/3 neurons can result in measurable biases in perceptual judgements of bilateral whisker input. This is consistent with several recent barrel cortex studies showing that stimulation of single neurons^33,40^, and small neural ensembles^35,102^, can evoke perceptual effects. Surprisingly, we found that perceptual bias was significantly predicted by the number of non-coding neurons activated by targeted photostimulation, challenging the consensus that decision-making processes read out activity from highly-tuned stimulus-responsive neurons^103^. Moreover, non-coding target neurons did not show sensory-evoked activity despite prominent increases in motor activity during task performance. This is intriguing, as recent studies have demonstrated that non-instructed movements evoked during sensorimotor behavior can profoundly influence the activity of cortical neurons^104,105^.

In contrast with recent studies involving the selective optical stimulation of functionally tuned neurons during behavior^41,45,62,102^, we did not find an effect of target ensemble tuning on task performance. This is likely because target group whisker-selectivity was relatively weak compared with previous studies, as we predominantly stimulated whisker non-coding neurons in the FOV. We anticipate this is because finding neurons with strong responses to both whisker input and 2P stimulation in the FOV was rare given that we imaged/stimulated and limited sample of neurons within a single plane. In future, this limitation could be resolved by optically interrogating larger^94^ and/or volumetric FOVs^44,106^. Thus, while further work is needed to investigate the differential contributions of intrahemispheric contra- and ipsi-coding neurons to task performance, we provide strong evidence that stimulus non-coding neurons can be recruited to enhance sensory perception.

However, it remains to be established how optogenetic recruitment of stimulus non-coding neurons would impact performance in other perceptual contexts. Recent work in visual cortex suggests that stimulating non-coding neurons can even impair performance by suppressing the perceptually relevant cells in the local circuit^45^, suggesting that the ability of non-coding neurons to influence perception may depend on task design or brain area. In our experiments, the dynamics of interhemispheric competition might engage a distinct decision-making process that emphasises rapid decoding of intrahemispheric spiking across a wider pool of neurons in barrel cortex in a winner-take-all scenario. Under these perceptually ambiguous conditions, any additional intrahemispheric spikes may thus help reach a decision threshold, which could also reflect increased capacity for the photostimulated hemisphere to suppress the contralateral hemisphere^78,79^, compensate for cortical adaptation to high-frequency whisker stimulation^107^ or enhance sensory responses in deeper cortical layers^108,109^.

### A reserve pool of silent neurons can be recruited to enhance stimulus perception

The observation of large fractions of stimulus non-responsive neurons in cortex has long been intriguing^21,46,47^. Non-responsive neurons could be deliberately suppressed by intrinsic or synaptic mechanisms to constrain excessive cortical excitation, reduce coding redundancy, and/or increase energy efficiency via sparse coding^50,110–112^. Alternatively, silent neurons may simply reflect structurally or functionally immature neurons that are not fully integrated into the circuit. Our targeted photostimulation results demonstrate that under our task conditions, non-coding neurons able to contribute to percept generation but are suppressed by strong network inhibition during task-related whisker processing. This is consistent with intracellular recordings showing that many L2/3 neurons are initially depolarized during whisker stimulation but then are rapidly hyperpolarized by subsequent cortical inhibition^7,19^. Our results suggest that releasing these neurons from inhibitory suppression, as we achieved through patterned optogenetic stimulation in our study, but which could be achieved endogenously via disinhibitory VIP+ interneuron circuits^113^ and cortical plasticity^20,49^, can provide a simple mechanism for enhancing intra-cortical sensory signals. Our finding that ‘silent’ neurons can contribute information to perceptual decisions also adds causal support to emerging evidence showing that neurons in sensory cortex without classical stimulus-driven responses can contribute to neural processing^114–116^. These results have important implications for designing novel therapeutic optical brain-machine interfaces and optogenetic therapies^117^, since it suggests that it may not be necessary to precisely target optogenetic interventions to specific functionally defined pools of neurons. Rather, activation of a relatively small pool of pyramidal neurons regardless of functional identity could be sufficient to enhance sensory coding and restore some basic perceptual functions.

## Acknowledgements

We thank Selmaan Chettih and Christopher Harvey for developing and sharing the somatically-restricted C1V1 opsin; Soyon Chun for mouse breeding; members of the UCL Biological Service Unit for animal care and husbandry; Hamish Forrest for behavioral training during pilot experiments; and Bruker Corporation for technical support. We also thank Ann Duan, Sonja Hofer and Miguel Maravall for useful discussion. This work was supported by grants from the Wellcome Trust (PRF 201225 and 224688), ERC (AdG 695709), MRC (MR/T022922/1) and the BBSRC (BB/N009835/1).

## STAR ★ Methods

### RESOURCE AVAILABILITY

#### Lead contact

Further information should be directed to Michael Häusser.

#### Materials availability

This study did not generate any new reagents or materials.

#### Data and code availability

Data and code will be made available upon reasonable request to the lead contact.

### EXPERIMENTAL MODEL AND SUBJECT DETAILS

All experimental procedures were carried out under license from the UK Home Office in accordance with the UK Animals (Scientific Procedures) Act (1986). Wild-type adult female mice (*Mus musculus*, C57BL/6, P35 - 42 on day of surgery) were used for all behavioral, imaging and photostimulation experiments. PV-Cre mice (Jax 008069) were used for optogenetic photoinhibition experiments. Mice were single-housed in individually ventilated cages (IVCs) equipped with environmental enrichment to avoid whisker barbering when group-housed. Behavioral training usually took place for 2-3 weeks, and then experiments were conducted for 2-3 weeks following task learning.

### METHOD DETAILS

#### Surgical procedures

Mice were implanted with a headplate, injected with virus and installed with a cranial imaging window in a single surgery session. Mice were first anaesthetised with isoflurane (5% induction, 1.5% maintenance) and injected subcutaneously with an analgesic (Carprieve). The scalp was shaved with clippers then cleaned with iodine and a physiological *in vivo* external (IVE) solution (150 mM NaCl, 2.5 mM KCl, 10 mM HEPES, 2 mM CaCl_2_, 1 mM MgCl_2_) using sterile swabs (Sugi, Kettenbach). Mice were then fixed in a cranial stereotaxic frame and placed on a heat mat maintained at 37°C. Lidocaine was applied topically to the scalp before incision. Scalp was then removed bilaterally with surgical scissors revealing the dorsal surface of the skull. An aluminium headplate with a 7 mm diameter circular imaging well was fixed over the right hemisphere with dental cement (Super-Bond C&B, Sun-Medical).

A craniotomy (4 mm diameter) was made in the centre of the headplate well over S1 (2 mm anterior, 3.5 mm lateral from bregma) with a dental drill (NSK UK Ltd.), and the dura was removed using fine tweezers. Virus was front-loaded into a bevelled micropipette and injected at cortical depth of 300 µm at 0.1 µl/min (total volume 0.5 – 1 µl) using a calibrated oil-filled hydraulic injection system (Harvard apparatus). For one-photon optogenetic experiments, mice were co-injected with GCaMP6s (AAV1-hSyn-GCaMP6s-WPRE-SV40) and C1V1 (AAVdj-CaMKIIa-C1V1(E162T)-TS-P2A-mCherry-WPRE) in a 1:10 ratio. For two-photon imaging and targeted two-photon photostimulation experiments, mice were co-injected with GCaMP6s (AAV1-hSyn-GCaMP6s-WPRE-SV40) and somatically-targeted C1V1 (st-C1V1; AAV2/9-CaMKII-C1V1(t/t)-mScarlett-Kv2.1). Stock st-C1V1 was diluted 1:10 in virus buffer, and GCaMP6s was added to the dilution mixture in a ratio of 1:10 GCaMP:st-C1V1. For PV-Cre mice, flexed-C1V1 (AAV-DJ-EF1a-DIO-C1V1(E162T)-TS-p2A-mCherry) was injected with GCaMP6s in a 1:10 GCaMP:C1V1 ratio. Opsin-negative control mice were injected with GCaMP6s diluted 1:10 in virus buffer. The needle was retracted slowly 5 minutes after the injection was completed and a cranial window (made from a 3 mm circular glass cover slip glued onto a 4 mm circular glass cover slip with optical glue NOR-61, Norland Optical Adhesive) was press-fit into the craniotomy and sealed in place with Vetbond and dental cement. Following completion of the surgery, mice were placed in an incubated recovery chamber and monitored closely until normal locomotor activity resumed. Post-operative care included daily weight and health monitoring, and Carprieve administration in the home cage water supply for 4-5 days.

#### Whisker stimulation

Contralateral and ipsilateral whiskers were deflected along the anterior-posterior axis with two piezoelectric actuators (Physik Instrumente; PL127.11) mounted on NOGA articulated arms (RS 785-7869) and positioned ~5 mm from the base of the whisker pad on either side of the snout. For single-whisker stimulation, the C2 whiskers were threaded into glass-capillaries glued to the piezo actuators and deflected in the anterior direction with a single 50 ms square pulse. For the single-whisker task every 1-2 weeks mice were briefly anaesthetised to allow retrimming of the surrounding whiskers. Piezo deflection range was 0 - 1 mm from rest as calibrated with videography. For multi-whisker stimulation, whiskers were deflected with cardboard stimulus ‘paddles’ (2 × 3 cm). Multi-whisker stimulation consisted of a sinusoidal deflection for 500 ms at 20 Hz with uniform intensity. Stimulus intensity was modulated by varying the voltage signal amplitude used to drive the piezos. Analogue waveforms were generated using custom written MATLAB (MathWorks) scripts using the ‘Data Acquisition Toolbox’, and outputted to single channel piezo drivers (Noliac, NDR6110) via a National Instruments card (USB-6351).

#### Behavioral training

We developed a bilateral decision-making task where head-fixed mice discriminated bilateral whisker input and reported decisions by licking left or right. To motivate task engagement, access to cage water was removed, and water (~1 ml per day, 5% sucrose solution) was received through daily behavioral training. Body weight was maintained between 80 - 90% for the duration of training, which typically lasted 1-2 months. Daily health checks were performed to identify signs of poor health and excessive dehydration, and supplementary water was provided if necessary. Prior to training, mice were first habituated to experimenter handling and allowed to freely explore the head-fixation platform and tube. During head-fixation habituation, mice were trained to tolerate periods of head-restraint of increasing duration, and water rewards were delivered through the dual lickport to encourage pro-active directional licking.

We counterbalanced the spatial stimulus-response contingency across different mice. Mice were assigned to either the symmetric (e.g. stimulus left, lick left) or asymmetric (e.g. stimulus left, lick right) task contingency and were not trained to switch between the two. To ensure reliable learning of the task framework, mice were initially trained on contra and ipsi unilateral stimuli (i.e., no distractor stimuli) of 100% intensity. Stimuli were delivered in a randomized order, with a maximum of 3 consecutive trials of the same type in a row. To promote associative learning of the stimulus-response contingency, during the first 2-3 sessions the target lick response was prompted with an automatic reward (‘autoreward’), which was unconditionally triggered 750 ms after the stimulus. Following 2-3 days of autoreward instruction, subsequent sessions had the autoreward feature disabled, requiring mice to respond correctly to obtain water rewards. Autoreward was occasionally and transiently re-enabled if mice displayed a strong response bias that persisted across multiple sessions. Correct trials were rewarded with a 5 µl water droplet (5 % sucrose solution), incorrect trials incurred a 5 s timeout while miss trials were not penalised.

For the delayed-response task version, following whisker stimulation (0.5 s), mice withheld licking and were instructed to report their decision after a ~1.5 second delay (from stimulus onset) via an auditory go cue (piezo buzzer, 200 ms). To enforce this delay period, the lickports were mounted on a linear motor (P16 Mini Linear Actuator, Actuonix), which was driven via a linear actuator control board (LAC board, Actuonix). The lickports were moved in and out on each trial using a square driving signal generated in MATLAB (2 s duration, duty cycle 20%). The lickport motor took 170 ms to extend (a translation of 1 cm). In imaging and photostimulation experimental sessions, a post-hoc video-based measure of licking was derived to quantify early lick responses.

Following learning of the basic task framework, mice were transitioned to intensity discrimination training. Simultaneous bilateral deflections were delivered to the whiskers, and the difference in deflection intensity cued the target behavioral response in line with the learned task contingency. In discrimination sessions incorrect choices were not punished with negative reinforcement (beyond the omission of reward) to encourage mice to respond across all trial regardless of discrimination difficulty, and threshold stimulus trials were rewarded with 50% probability.

The behavioral task configuration was designed and controlled using PyBehavior (https://github.com/llerussell/PyBehavior). Behavioral training took place in custom built training boxes lined with sound attenuating foam. A dual lickport was constructed using syringe needles (manually filed down to a dull point) held in a plastic block (distance between syringe tips 5 mm), and a dual lick-o-meter circuit was used to detect licks electrically. Each lickport was gravity-fed with water through plastic tubing, which was gated by a solenoid pinch valve (225PNC1-11, NResearch). Reward volume was 5 µl (5 % sucrose solution). A set of USB computer speakers provided constant background white noise to mask sensory cues (e.g., piezo movement) and external noise. All behavioral training and experiments were performed in the dark.

#### Muscimol silencing

To allow pharmacological access to cortex, we installed cranial windows with a laser cut hole through the middle (500 µm diameter, Laser Micromachining, UK) plugged with silicone (Kwik-Sil) and centred over barrel cortex in 4 mice. The dura was removed before the cranial window was installed. Prior to the experiment, the silicone plug was removed from the laser-cut hole in cranial window, permitting access to cortex. Muscimol (powder dissolved in IVE; 100 nl at 5 µg/µl; Sigma-Aldrich) was manually pipetted onto the cranial window hole using a Gilson pipetter and allowed to diffuse into barrel cortex for 20 minutes. Task performance was assessed before and after muscimol application, as well as 24 hours after application in a recovery session.

#### Widefield calcium imaging

Widefield GCaMP fluorescence imaging was performed to localize barrel cortex (and individual whisker barrels) through the cranial window 2-3 weeks following surgery. Cortex was illuminated with a blue LED (Thorlabs) focused onto the cortical surface through a 4x/0.1-NA air objective (Olympus). GCaMP6s fluorescence was passed through a GFP excitation filter (Thorlabs) and detected using a CMOS camera (ORCA-Flash 4.0, Hamamatsu) via the objective (FOV ~1.5 × 1.5 mm). Contralateral whiskers were deflected at 10 Hz for 0.5 seconds during imaging. Stimulus-triggered average images were normalized to a 3 s pre-stimulus baseline window. Widefield imaging was performed before every imaging and photostimulation experiment at 10 fps, 15-20 stimulus repeats were delivered passively (i.e. not during task performance) to reduce licking/reward signal contamination.

#### One-photon optogenetics

All one-photon photostimulation experiments were performed with an LED (595 nm, Thorlabs M595L3), mounted in the imaging light path above the two-photon objective. One-photon stimulation was delivered through the two-photon objective, which was positioned over the C2 whisker barrel in right hemisphere barrel cortex. LED power (mW) and triggering was controlled via an LED driver T-Cube (Thorlabs). Power was calibrated with a power meter (PM100A, Thorlabs). For the optogenetic substitution experiments presented in Figure 2A-C (C1V1 in pyramidal neurons), the LED stimulus was a square pulse, 50 ms, power range: 0-50 mW. For photostimulation biasing experiments in Figure 2F (C1V1 in pyramidal neurons) the LED stimulus was a single square pulse (50 ms, 1 mW). For optogenetic photoinhibition experiments in Figure 2G (C1V1 expressed in PV-interneurons), the LED stimulus was a 500 ms continuous square pulse at 10 mW. Trials with optogenetic stimuli were not rewarded to avoid reinforcing behavioral responses and were interleaved with a high proportion of whisker trials to maintain task engagement. The only exception to this is the experiments presented in Figure S4D, where mice were intentionally trained to report optogenetic stimulation.

#### All-optical system design

We used an ‘all-optical’ experimental strategy^53–55,118^ to investigate the impact of sparse activation of L2/3 cortical ensembles on perception and local circuit physiology. All-optical experiments were performed on a customized dual beam path ‘all-optical’ microscope described previously^53^. A photostimulation light path coupled to a spatial light modulator (SLM) enabled spatially precise two-photon photostimulation (PS), while a second imaging light path provided optical readout of activity via 2P calcium imaging. Imaging data were acquired on a resonant scanning system (Bruker Corporation) at 30 Hz with 512 × 512-pixel resolution (FOV 470 × 470 µm). GCaMP6s was excited using a Chameleon Ultra II laser (Coherent) at 920 nm, which was raster scanned across the FOV. Static C1V1-mCherry and st-C1V1-mScarlet images were acquired at 765 nm. Imaging power under the objective was 50 mW. A 25x/0.95-NA water-immersion objective (Leica) was used for all experiments. Two-photon excitation of soma-targeted (st) C1V1 expressing neurons was performed with a femto-second pulsed laser at 1030 nm (Satsuma, Amplitude Systèmes, 2 MHz repetition rate, average output 20 W, pulse width 280 fs) coupled to a spatial light modulator (SLM, 7.68 × 7.68 mm active area, 512 × 512 pixels, Meadowlark Optics/Boulder Nonlinear Systems). SLM phase masks were generated using the Gerchberg-Saxton algorithm^119^ and displayed on the SLM display using control software (Blink, Meadowlark). SLM targeting precision was ensured using calibration routines that mapped SLM pixel space onto 2P pixel space via an affine transformation^53,54^. SLM-generated beamlets were simultaneously spiral scanned (three rotations, 15 ms duration per spiral, 15 µm diameter at 20 Hz for 500 ms) over somata with a pair of galvanometer mirrors. Photostimulation power was calibrated at 6 mW per cell. Power was modulated via an acoustic optical modulator (AOM), and calibrated with a power meter (PM100A, Thorlabs). Recordings were performed in 7 symmetric-trained mice (30 sessions, 4.3 ± 2.0 sessions per mouse) and 6 asymmetric-trained mice (22 sessions, 3.7 ± 1.8 sessions per mouse).

#### Photostimulation response mapping

Due to variable levels of GCaMP6s and st-C1V1 co-expression, not all cells in a FOV are addressable for all-optical interrogation. In order to identify cells with both functional opsin and indicator, we used a flexible photostimulation mapping procedure designed to enable rapid tests of the photostimulation responsivity of each cell in the FOV (Near automatic photoactivation response mapping: ‘*NAPARM’*^44,54^). Static expression images of GCaMP and st-C1V1 were loaded into NAPARM’s user GUI, and 350 pixel targets (corresponding to the xy position of cell somata in the 2P FOV) were selected semi-automatically based on local intensity maxima. The 350 photostimulation target sites were clustered into 7 target groups of 50 cells using ek-means, and a phase-mask and a set of photostimulation galvanometer position coordinates for each cluster was generated. Sequential photostimulation of each target group was performed by the photostimulation module of the all-optical system and the SLM software and was triggered using synchronisation software (PackIO; https://github.com/apacker83/PackIO). 15 photostimulation repeats of the stimulation sequence (stim pattern 1 through to 7) were performed (total of 105 photostimulation events per experiment). Photostimulation parameters were 15 ms spiral, 15 μm diameter, 20 Hz, 500 ms duration, 10 reps, 5 s ITI. Photostimulation responsive neurons were identified by comparing baseline ΔF/F responses (mean in a 1 s pre-stimulus window) to evoked ΔF/F responses (mean of 0.5 to 1 s post-stimulus) with a one-tailed Wilcoxon ranked-sum test (P < 0.05 = responsive cell).

#### Two-photon photostimulation group selection

In each experiment, two stimulation target groups were selected as the photostimulation-responsive neurons with the top 30 and bottom 30 whisker stimulus-selectivity scores, which were characterized based on a short behavioral imaging session.

### QUANTIFICATION AND STATISTICAL ANALYSIS

#### Behavioral metrics

To track learning across sessions, we quantified the overall fraction of correct trial (P(Correct)) across contra and ipsi unilateral trials excluding miss trials. For quantification of discrimination accuracy, we analysed trials where mice reported a decision and excluded miss trials. For perceptual choices, we quantified the fraction of trials where mice reported the contra whisker ‘P(Report contra whisker)’. For motor (lickport) choices, we quantified the fraction of trials where mice licked the contra lickport ‘P(Lick contra lickport)’. We fit psychometric data using the sigm_fit function in MATLAB (https://www.mathworks.com/matlabcentral/fileexchange/42641-sigm_fit). We used the fit to estimate the psychometric threshold stimulus (TS) as the corresponding stimulus condition that would result in 50% discrimination performance on the psychometric curve. For assessing the perceptual effect of optogenetic manipulation on TS trials discrimination, we quantified the difference in P(Report contra whisker) on control TS trials and opto+TS trials.

#### Two-photon calcium imaging data

All two-photon imaging data were motion-corrected and segmented into somatic and neuropil fluorescence traces using the Python release of Suite2p^120^ (https://github.com/MouseLand/suite2p). Manual curation was performed to discard ROIs with non-somatic shapes (e.g. dendritic/axonal processes) or filled nuclear fluorescence. Neuropil subtraction was performed across cells by subtracting the neuropil signal from the somatic signal. Prior to subtraction, the neuropil signal was scaled by a coefficient (ranging from 0 – 1), which was based on an estimation of neuropil signal contamination in the somatic signal using robust linear regression^121^.

For each neuron and each trial, the neuropil-subtracted fluorescence signal was extracted 1 second pre-stimulus to 6 seconds post-stimulus (210 frames in total). Each trial trace was normalised using a ΔF/F calculation, where F was the average fluorescence signal across the 1 second pre-stim baseline (30 imaging frames). The evoked response on a given trial was subsequently calculated as the average ΔF/F signal 0.5–1 second post-stimulus (15 imaging frames). This analysis window avoids both sensory and photostimulus presentation (which last 0.5 s) and avoids licking. Thus, the imaging response window is optimized to measure the stimulus-evoked signal prior to contamination from response/reward related activity.

#### Identifying neurons with significant trial responses

Neurons with significant trial-evoked responses were identified by comparing the evoked ΔF/F response (during the delay epoch 500 – 1000 ms after stimulus onset) distributions across different trial conditions using a one-tailed Wilcoxon rank-sum test and a P < 0.05 threshold. For example, neurons activated by photostimulation (PS) were identified by comparing PS trial to Catch trial response distributions. Likewise, neurons activated by photostimulation on top of sensory input (TS+PS) were identified by comparing responses on TS trials to TS+PS trials. Neurons significantly selective for contra vs ipsi were defined by comparing trial-wise evoked responses across contra and ipsi trials.

#### Pixel-wise analysis of imaging data

Pixel-wise analysis was performed on the raw calcium imaging data to corroborate trace-based analyses. This was important for confirming that Suite2p-detected responses (e.g. stimulus-selective / photostimulation activated) reliably reflected measurable and visible changes in somatic fluorescence (as opposed to neuropil contamination or passing axonal/dendritic processes). For each trial, registered imaging frames were normalised (ΔF/F, as described above) to a 1 second pre-stimulus baseline, and STA response was assessed as the average response (500-1000 ms) post stimulus. Pixelwise STAs ‘stamps’ for individual cells were generated by sampling a mini-FOV 50 × 50 μm with each cell centred in the stamp at pixel location (25,25).

#### Receiver operating characteristic (ROC) analysis

We used ROC analysis (MATLAB’s perfcurve function) to assign a stimulus-selectivity score to each neuron according to how well an ideal observer could decode trial-type (Contra vs Ipsi) from the evoked response. Stimulus selectivity was defined as the area under the ROC performance curve (AUC). Neurons with stimulus selectivity > 0.5 were selective for Contra trials, and neurons < 0.5 were selective for ipsi trials. We used the same procedure to quantify each neurons correlation with behavioral choices on threshold trials by calculating action probability (comparison of 21.7 ± 15.7 ‘lick contra lickport’ and 24 ± 14.9 ‘lick ipsi lickport’ trials), choice probability (comparison of 21.6 ± 14.6 ‘report contra whisker’ and 24.1 ± 15.3 ‘report ipsi whisker’ trials) and detect probability (comparing lick vs no lick trials).

#### Behavioral videography

Behavioral videography was acquired during two-photon imaging and targeted two-photon photostimulation experiments using two infra-red (IR) sensitive CMOS cameras (DCC3240M Thorlabs) under infra-red LED illumination. One camera positioned face-on to the mouse (‘FaceCam’), recorded orofacial movements (licking and whisking) with a framerate of 75 – 100 fps. A second camera (‘BodyCam’) recorded body movements at 20 fps. Acquisition parameters (FOV size, frame rate etc.) were configured using ThorCam (Thorlabs).

To analyse whisker pad and body movements, we used an ROI-based procedure to extract movement (in arbitrary units) in the video recordings. Rectangular ROIs were positioned over the contra and ipsi whisker pads and across the body. We computed the correlation coefficient ‘r’ between frame-to-frame pixel intensities as a measure of frame-to-frame similarity. Consecutive frames without any animal movement (e.g. no whisking) will have a ‘r’ close to 1. On the other hand, frames with a lot of movement will have a low ‘r’, as pixel intensities for a given pixel location will vary from moment to moment. Frame to frame whisking and body activity was therefore summarised as ‘1 – r’, normalised to the maximum value.

We used DeepLabCut^122^ (DLC; https://github.com/DeepLabCut/DeepLabCut) to quantify licking responses that occurred outside of the response window (whilst the motorised lickport was retracted). A single ‘global’ DLC model was trained using FaceCam video frames from across a sample of experiments. The model was trained using an equal proportion of licking / no licking frames (selected manually), with 10 ‘nodes’ manually-labelled on the tongue (9 around the edge, 1 in the centre) on ~200 frames. On ‘test’ data, DLC tongue labels were assessed visually, and the model retrained if tracking accuracy was deemed poor. A binary video measure of licking (tongue present / not present) was calculated by assessing if 7 or more tongue labels were present in a given frame each with a ‘likelihood’ of ≥ 0.7.

#### Control trial resampling

To test whether the relationship between photostimulation target count and perceptual bias reflected an underlying correlation between the number of active neurons and perceptual choice we performed a shuffle test based on resampling non-photostimulation trials. For each experiment, we sub-sampled (with re-sampling) control TS trials, to match the corresponding number of photostimulation trials. This was possible as TS trials were delivered with twice the frequency compared to each of the TS+PS trial conditions. We then compared the choice deviation ‘ΔP(Report contra whisker)’ of this sub-sample to the average of the control TS trials, and summed the number of photostimulation target neurons that were identified as being significantly activated. We did this twice for each experiment to generate a ‘fake’ TS+PS dataset, and then assessed the correlation between choice bias and photostimulation target activity. We repeated this procedure 10000 times to build up a distribution of correlation coefficients against which we could compare the true photostimulated trials with.

#### Statistical analyses

Unless otherwise stated all paired statistical tests used were Wilcoxon signed-rank tests. For unpaired comparisons Wilcoxon rank-sum tests were used. A linear regression model (Matlab ‘*fitlm’*) was used to predict photostimulation-evoked choice bias using various photostimulation and behavioral parameters. All inputs to the linear regression model were first z-scored. Scatter plots show least-squares lines, and correlations were assessed using the Matlab function ‘*corrcoef’.* All data are reported as the mean and SEM (unless otherwise stated).

