## Supplementary figures (S1 - S13) for "A latent pool of neurons silenced by sensory-evoked inhibition can be recruited to enhance perception"

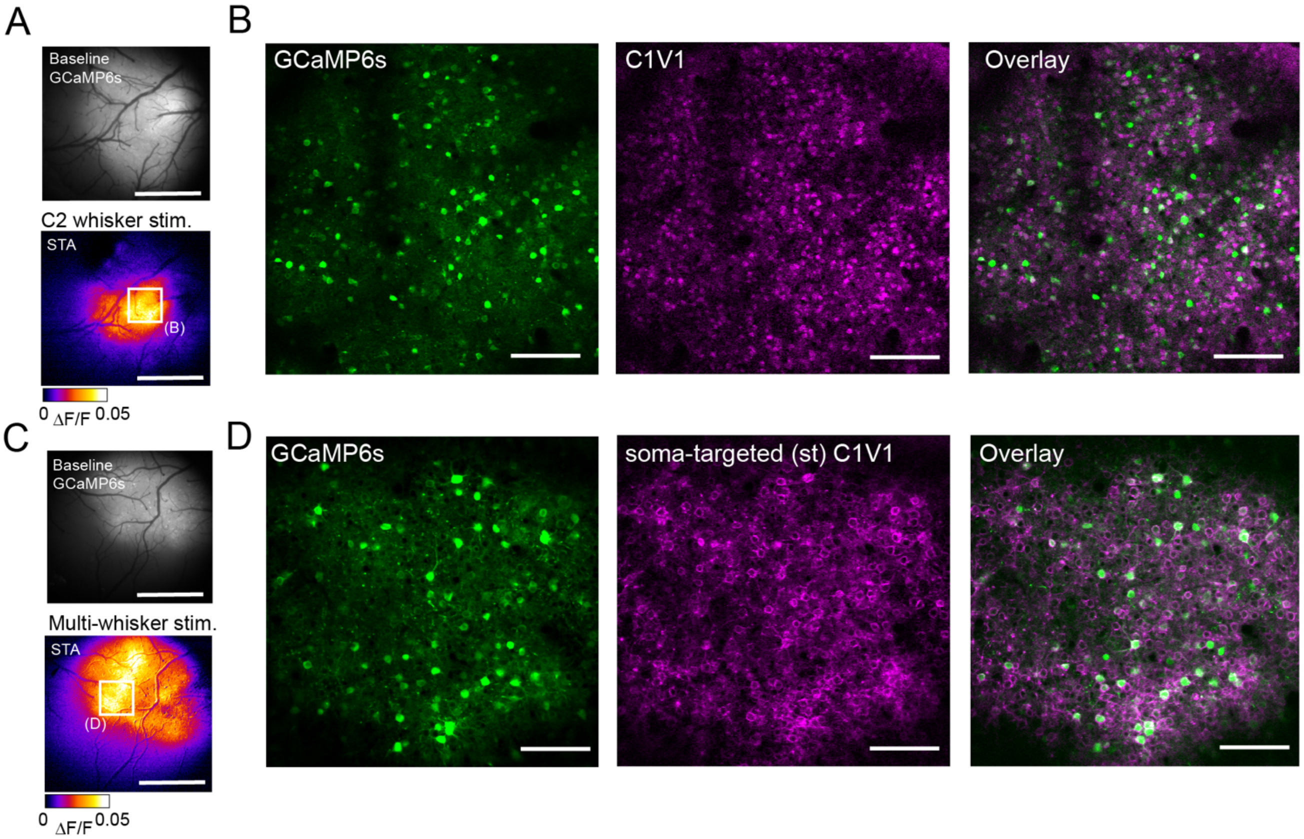


**Figure S1. Co-expression of GCaMP6s and C1V1 in barrel cortex.**

**(A)** Localisation of the C2 whisker barrel using widefield fluorescence imaging. Top image shows widefield baseline GCaMP6s fluorescence recorded through the cranial window. Bottom image shows the stimulus-triggered average (STA) fluorescence response following stimulation of the contralateral C2 whisker. The STA image shows the average ΔF/F response 0.5 - 1 s post stimulus. The white square indicates the location of the 2P field of view shown in panel (B). **(B)** 2P images of GCaMP6s (left, 920 nm), C1V1-mCherry (middle, 765 nm) co-expression in L2/3 barrel cortex. The right image shows the overlay. **(C)** Same as (A) but showing the widefield cortical ΔF/F response 0.5 - 1 s after the simultaneous stimulation of multiple contralateral whiskers in a different example mouse. The white box indicates the FOV location in (D). **(D)** 2P images of GCaMP6s (left, 920 nm), st-C1V1-mScarlet (right, 765 nm) and the overlay (right) expressing L2/3 neurons in barrel cortex. Images in (D) correspond to the FOV depicted in Figure 3D. Scale bar for widefield images = 1 mm. Scale bar for 2P images = 100 μm.

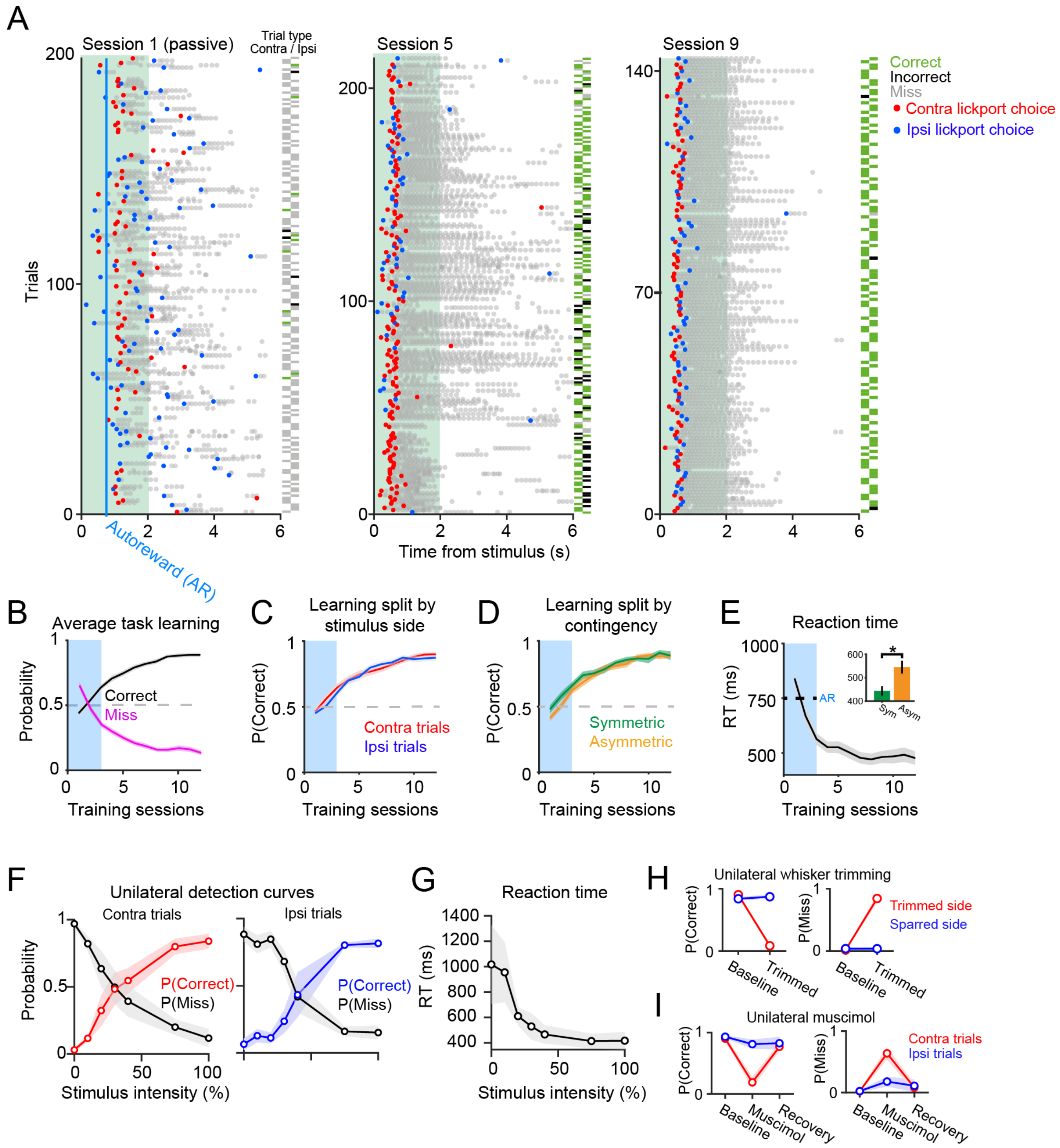


**Figure S2. Task learning and behavioural control experiments**

**(A)** Behavioural performance plots from an example animal in 3 different sessions across learning. The vertical blue line indicates the time of the autoreward (750 ms) - which was used to shape learning during early sessions. Each row corresponds to a trial, and each marker shows a lick. The first post-stimulus lick is coloured red/blue according to contra/ipsi lickport choice. Trial-type and trial outcome are indicated by the location and colour of the rectangular ticks on the right side of the session plot. **(B)** Average performance curves. P(Correct) and P(Miss) rate are shown in black and magenta across sessions. Data show the mean ± SEM across 61 mice; 259 ± 79 trials per session. **(C)** Average performance split by contra (red) and ipsi (blue) trials across learning. **(D)** Average performance for symmetric-trained mice (green; n = 31 mice) and asymmetric-trained mice (orange; n = 30 mice) across learning. **(E)** Average reaction times for correct trials across learning. The inset shows quantification of mean RT across symmetric and asymmetric mice (on sessions with P(Correct) > 70%). P < 0.05; Wilcoxon rank sum test. **(F)** Average performance ('P(Correct)') and miss rate across different unilateral deflection intensities for contra (left) and ipsi (right) stimulus trials. **(G)** Average reaction times across different deflection intensities. Data in F and G are from 4 mice (1 session per mouse). **(H)** Task performance before and after unilateral whisker trimming. Whisker trimming experiments were performed in 8 mice on the final day of experiments/training. (I) Task performance before and 20 minutes after unilateral muscimol infusion in barrel cortex. Recovery performance was assessed 24 hours later. Muscimol experiments were performed in 4 mice (1 session per mouse) trained on the symmetric-contingency.

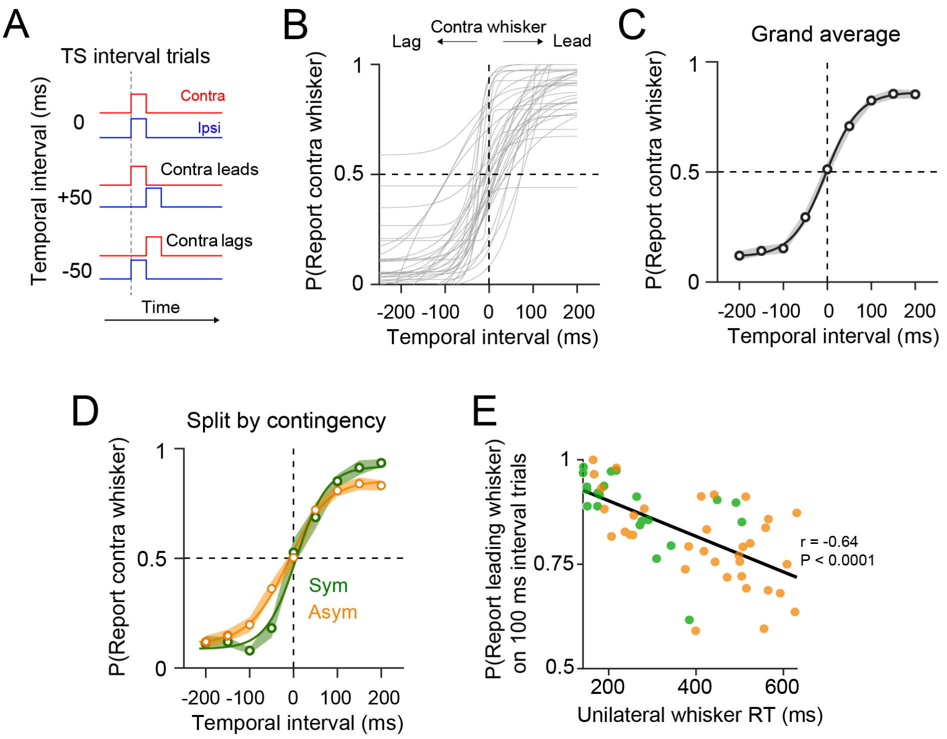


**Figure S3. Temporal discrimination of bilateral whisker stimulation**

**(A)** Schematic overview of TS interval trials. To probe temporal sensitivity to intervals in bilateral whisker stimulation mice received trials where the contra and ipsi component of the bilateral threshold stimulus (TS) were offset in time. TS-interval trials were not rewarded and were interleaved with a high-proportion of unilateral trials to maintain task engagement. **(B)** Choice tendency 'P(Report contra whisker)' is shown as a function of bilateral whisker temporal interval. Each line represents a psychometric curve from an individual training session (n = 58 sessions, 25 mice). **(C)** Average temporal discrimination curve generated from the data in (B). **(D)** Average temporal discrimination curves are shown for symmetric (green) and asymmetric (orange) trained mice. **(E)** Leading whisker bias on 100 ms interval trials is plotted against mean reaction times on unilateral whisker trials. Individual points show single sessions from symmetric (green) and asymmetric (orange) trained mice (23 sessions in 9 symmetric, 35 sessions in 16 asymmetric mice).

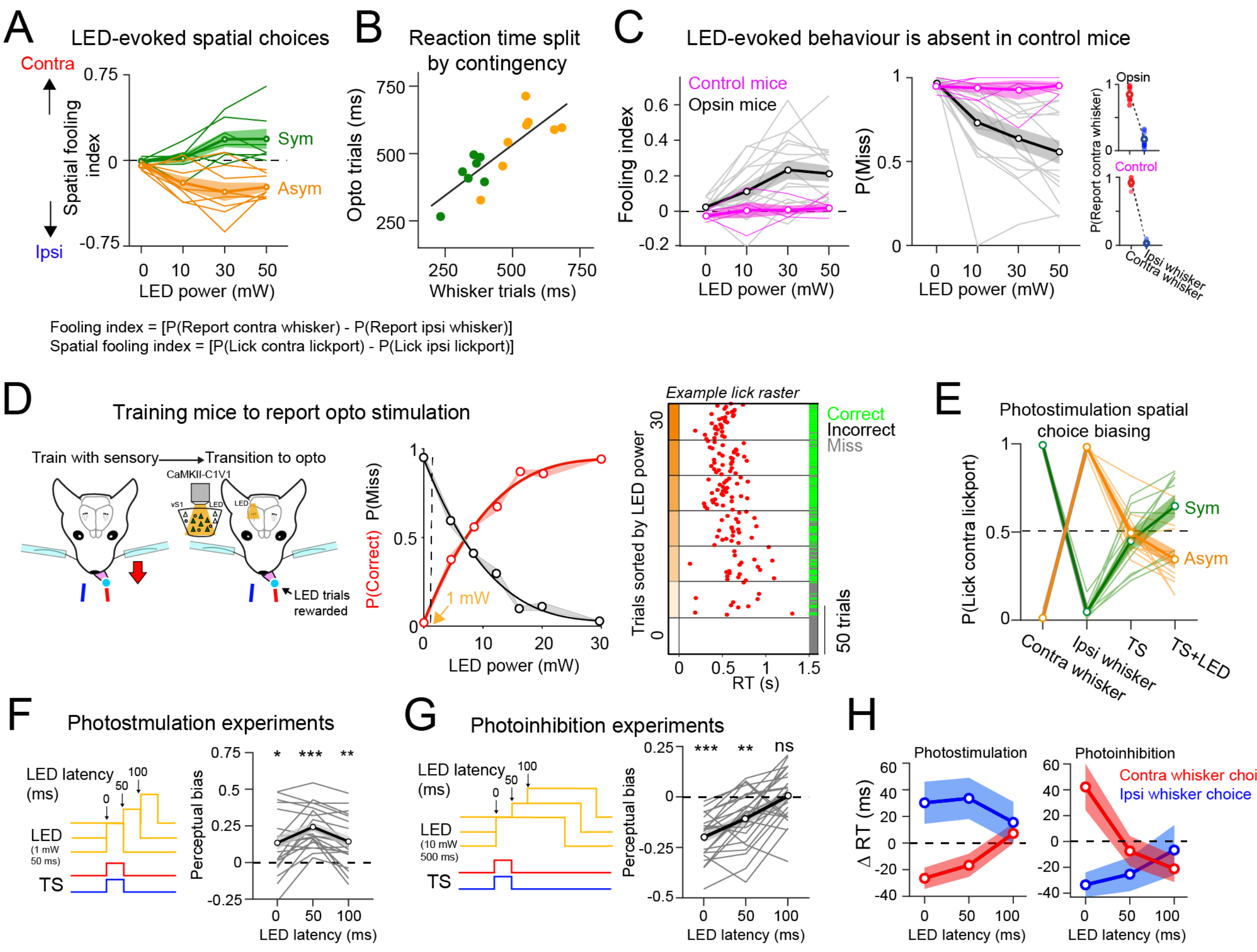


**Figure S4. Optogenetic manipulation of task performance: additional analyses and control experiments**

**(A)** Optogenetic 'substitution' experiments split by task contingency (n = 8 sessions in 4 symmetric-trained mice, and n = 8 sessions in 8 asymmetric-trained mice). **(B)** Average reaction time (RT) during optogenetic 'substitution' experiments showing symmetric (green) and asymmetric (orange) trained mice. **(C)** Optogenetic stimulation does not evoke perceptual responses in control mice (n = 5 sessions in 4 mice). Left: Fooling index for opsin expressing mice (grey) and control mice (magenta). Middle: Same as Left but showing P(Miss). Right: Summary of behavioural performance on whisker trials during the optogenetic experiment for opsin expressing mice (top) and control mice (bottom). **(D)** Perceptual detection of optogenetic stimulation. A subset of mice were trained on whisker trials, then transitioned to solely detecting LED stimulation. Licking responses on LED trials were rewarded, and so mice were motivated to detect LED stimulation. This experiment allowed us to identify a weak LED power condition (1 mW, indicated with the orange arrow) for optogenetic biasing experiments in Figure 2F. Data are from 2 mice (initially trained on the symmetric sensorimotor contingency). The lick raster plot shows an example optogenetic detection session. Contralateral lick reaction times are plotted as red circles. The LED power was modulated (0 – 30 mW, randomised) across different trials but sorted along the y- axis for display. **(E)** Spatial choice tendency during photostimulation-biasing sessions split by symmetric (green) and asymmetric (orange) trained mice. **(F)** Left: Schematic overview of photostimulation biasing experiments. C1V1-expressing excitatory neurons were optogenetically stimulated with LED input (50 ms pulse duration), which was triggered at 0, 50 and 100 ms relative to the whisker Threshold Stimulus (TS). Right: Perceptual bias on TS+LED trials is shown for different LED latencies. **(G)** Left: Schematic overview of cortical photoinhibition biasing experiments. Unilateral photoinhibition was performed by activating PV interneurons expressing C1V1 with an LED. Photoinhibition was triggered at 0, 50 and 100 ms relative to the TS. Right same as in (F) but for perceptual biasing on photoinhibition experiments across LED latency trials. **(H)** Average reaction time biasing on photostimulation (left) and photoinhibition (right) biasing sessions across LED latency trials. Statistical tests were Wilcoxon signed rank tests; n.s. P > 0.05; * P < 0.05; ** P < 0.01; *** P < 0.001.

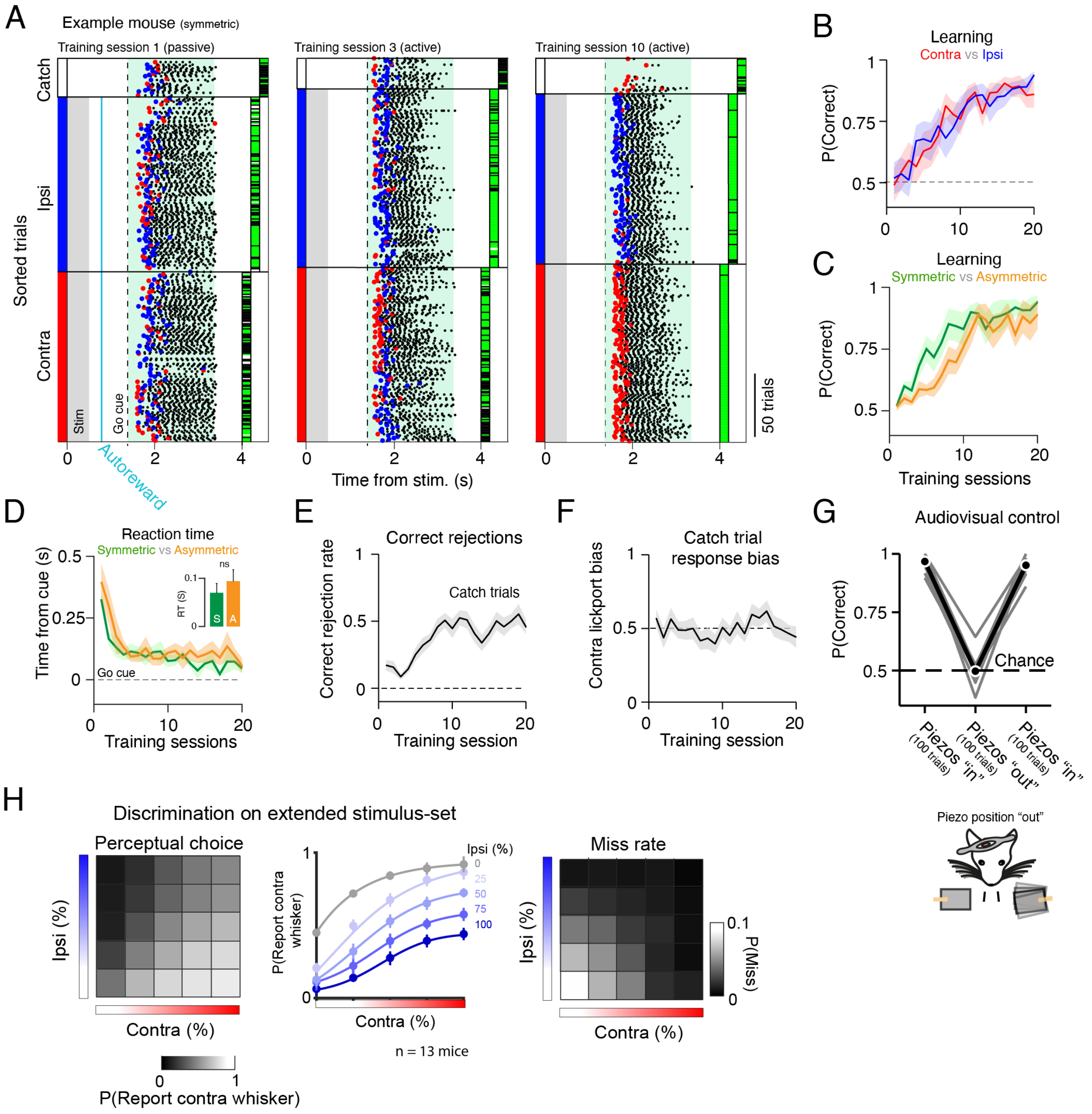


**Figure S5. Behavioural data across delayed-response task learning**

**(A)** Lick plots for an example symmetric-trained mouse across different stages of training. Trials are sorted along the y-axis according to stimulus type (contra, ipsi and catch). The blue vertical line in session 1 indicates the time of the programmed autoreward. Red/blue markers denote contra/ipsi lickport choice. The coloured ticks to the right of the axis denote correct (green), incorrect (black) and miss (white) trial outcomes. **(B)** Learning curves for contra (red) and ipsi (blue) trials. **(C)** Learning curves for symmetric (green, 7 mice) and asymmetric (orange, 6 mice) mice. **(D)** Average reaction times on correct trials across learning split by task contingency as in (C). Comparison was tested with a Wilcoxon rank sum test. n.s. P > 0.05. **(E)** Correct rejection rate on catch (no stimulus) trials across training. **(F)** Average response bias on catch trials across training. **(G)** Discrimination performance on unilateral trials was compared when the whisker deflector paddles were in contact with the whiskers ("in"), and when the paddles were positioned in an anterior position ("out"). This confirmed that mice do not sure audio-visual cues to solve the task. Individual grey lines show individual sessions. **(H)** Discrimination accuracy for bilateral discrimination sessions during training. During some training sessions, mice received 25 different stimulus combinations from a 5x5 stimulus matrix. Data in this figure are pooled across symmetric (7 mice) and asymmetric-trained (6 mice) mice.

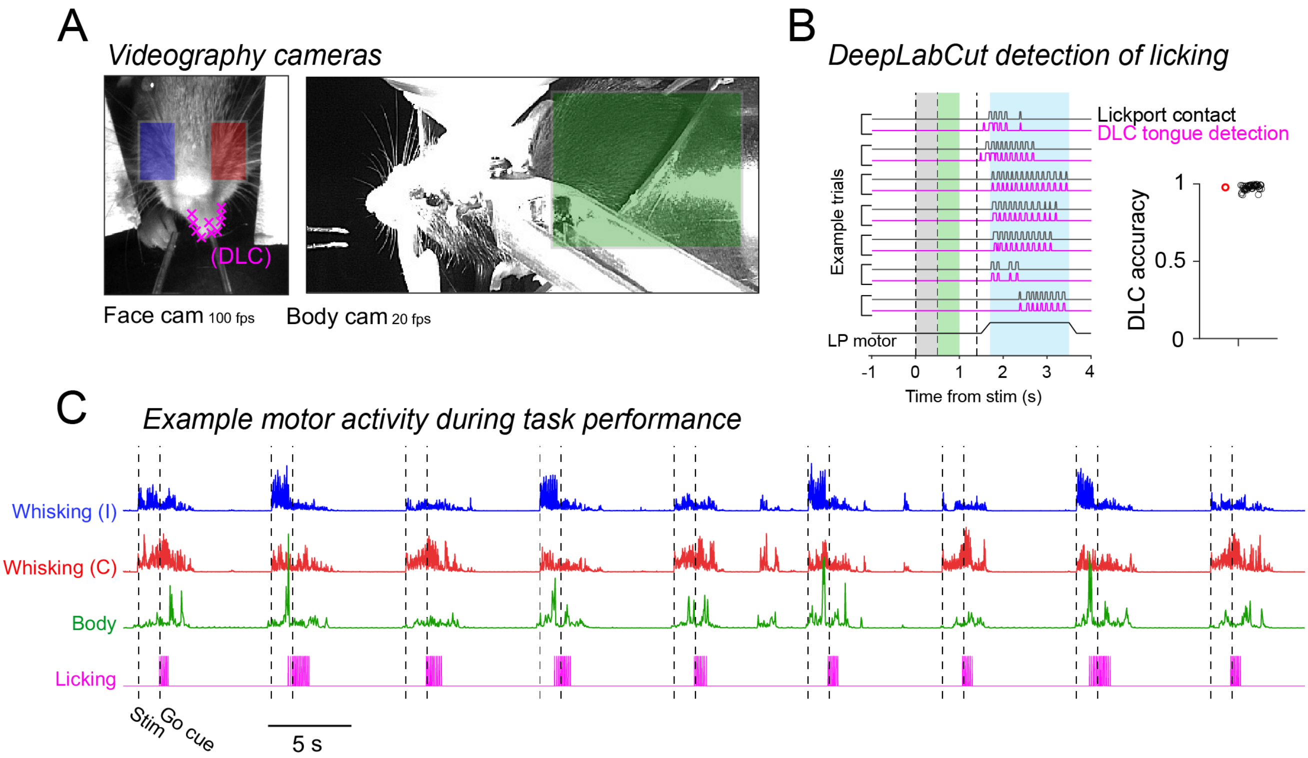


**Figure S6. Characterising movements during task performance using behavioural videography.**

**(A)** Example images show the standard view from 2 cameras, one set to record facial activity (left), and another set to record body movements (right). The coloured rectangular boxes indicate ROIs positioned to extract whisker pad movements (blue, red) and body movements (green) during task performance. The magenta 'x' markers indicate DeepLabCut tracking points tracing the outline of the tongue. **(B)** Comparing video-detected licks (magenta) with electrical lickport-detected licks (black) on 7 example trials. The summary plot shows quantification accuracy of video-detected licks compared with 'ground truth' lickport-detect licks. The red marker shows the overall average, while the black circles show the average performance for each session (52 session in 13 mice). On average, DeepLabCut performed with high accuracy (~98%; mean accuracy across 52 session) at detecting the tongue during licking behaviour. **(C)** Example whisking (red, blue), body movement (green) and licking (magenta) quantified using videography across a short section (9 trials) of continuous task performance.

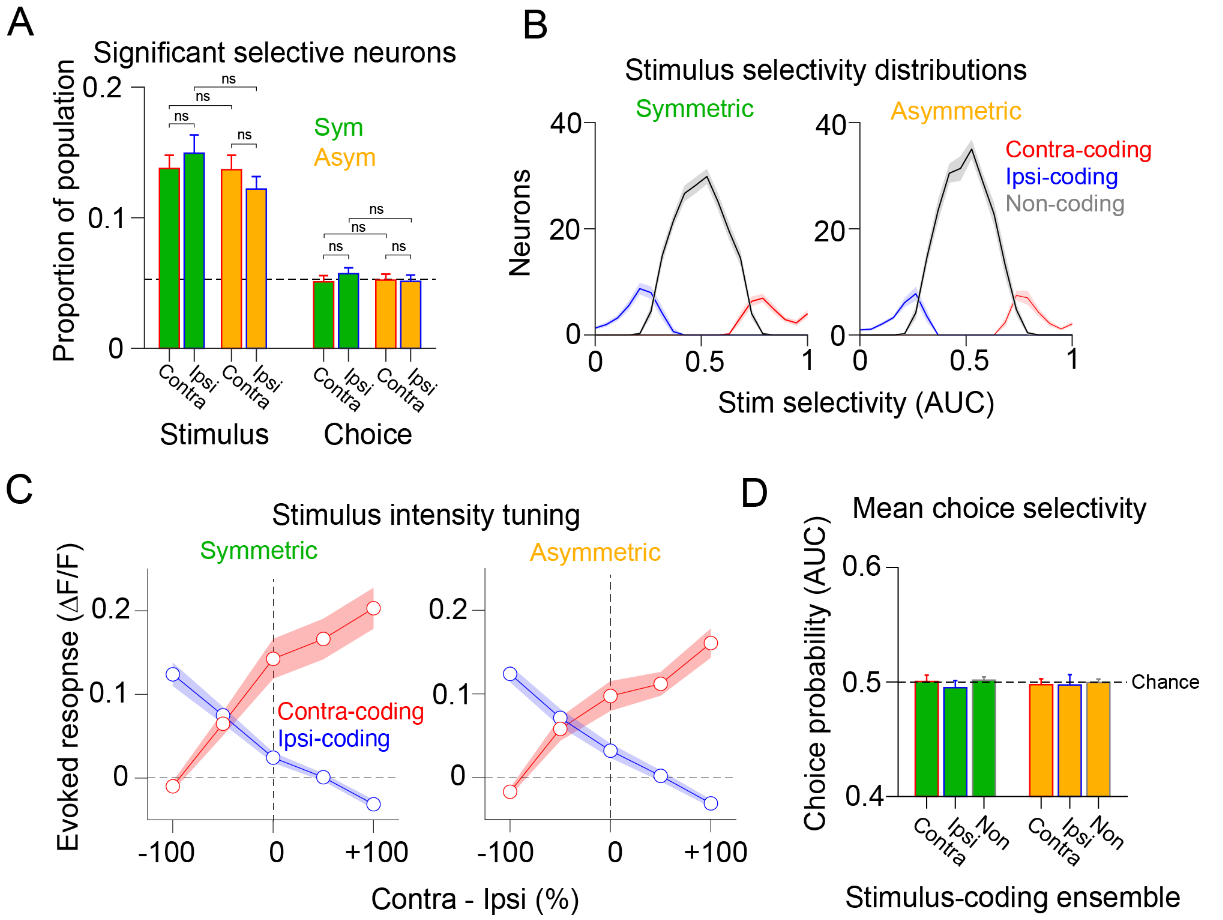


**Figure S7. Task-evoked neural activity is comparable across symmetric and asymmetric contingencies.**

**(A)** The fraction of statistically significant stimulus and choice-selective neurons for symmetric (green) and asymmetric-trained (orange) mice. **(B)** Average distribution of stimulus-selectivity scores for symmetric (left) and asymmetric (right) trained mice. **(C)** Mean evoked response size across trial-types in stimulus coding ensembles for symmetric (left) and asymmetric-trained (right) mice. **(D)** Mean choice probability scores across stimulus-coding and non-coding ensembles for symmetric (green) and asymmetric-trained (orange) mice. Data in this figure are from 30 sessions in 7 symmetric-trained mice and 22 sessions in 6 asymmetric-trained mice. Average numbers of neurons per FOV: Symmetric 253.4 ± 44.7, Asymmetric 260.7 ± 43.9. Average ensemble size: contra-coding sym 36.2 ± 16 neurons; ipsi-coding sym 38.4 ± 18.8; non-coding sym 189.1 ± 37; contra-coding asym 37.3 ± 15.7 neurons; ipsi-coding asym 35.2 ± 17.1; non-coding asym 209.3 ± 35 (mean ± std). Statistical tests within contingency groups were Wilcoxon signed rank tests. Statistical tests across contingency groups were Wilcoxon rank sum tests. n.s. $P > 0.05$.

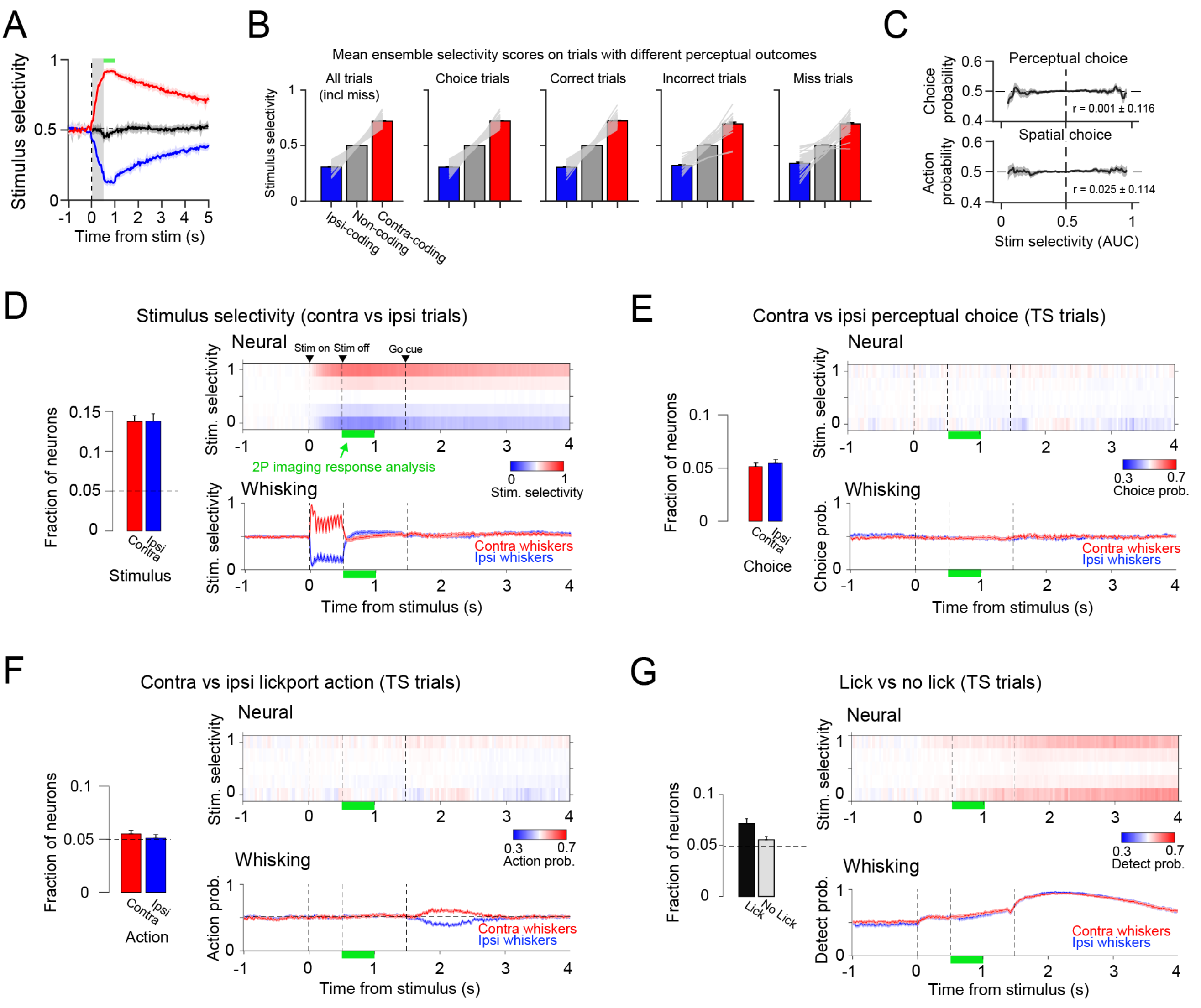


**Figure S8. Stimulus-coding neurons do not predict behavioural choice.**

**(A)** Average stimulus-selectivity in contra-coding (red), ipsi-coding (blue) and non-coding (black) ensembles across the trial. **(B)** Average selectivity scores (during the response analysis window indicated by the green bar in (A)) on contra vs ipsi trials in ensemble groups split by trials with different behavioural outcomes. Individual grey lines show individual sessions, the coloured bars show the average across sessions. **(C)** Top: Summary of the correlation between stimulus-selectivity scores and choice probability (on TS trials) across all neurons in the FOV. The plot shows average CP scores in neurons binned by stimulus-selectivity. The line and error bars indicate the mean and SEM across all sessions. The mean correlation strength (Pearson's corr) across sessions is indicated on the plot. The bottom plot shows the same but for Action probability scores. **(D)** Overview of neural and whisking decoding of contra vs ipsi stimulus type. Bar plot: the average fraction of significantly selective stimulus-coding neurons in the FOV. The horizontal dashed line at 0.05 indicates chance (false pos. rate based on a P < 0.05 statistical threshold for detecting significant responses). Neural (top; heatmap colour shows neuronal stimulus-selectivity across time (x-axis) across all neurons binned by stimulus-selectivity scores (y-axis) across all 52 sessions) and whisker (bottom; traces show analysis of whisking from the contra and ipsi side of the snout) decoding of contra vs ipsi whisker trials is shown across the trial epoch. **(E)** Bar plot: the average fraction of significantly selective whisker-choice coding neurons in the FOV. Neural (top) and whisker (bottom) decoding of perceptual choice is shown across the trial-epoch. **(F)** Bar plot: the average fraction of significantly selective lickport-choice coding neurons in the FOV. Neural (top) and whisker (bottom) decoding of lickport choice is shown across the trial-epoch. **(G)** Bar plot: the average fraction of significantly selective 'lick' vs 'no lick' coding neurons in the FOV. Neural (top) and whisker (bottom) decoding of 'lick' vs 'no lick' trials is shown across the trial-epoch. Data in this figure are from 52 sessions in 13 mice and show the average ± SEM across sessions.

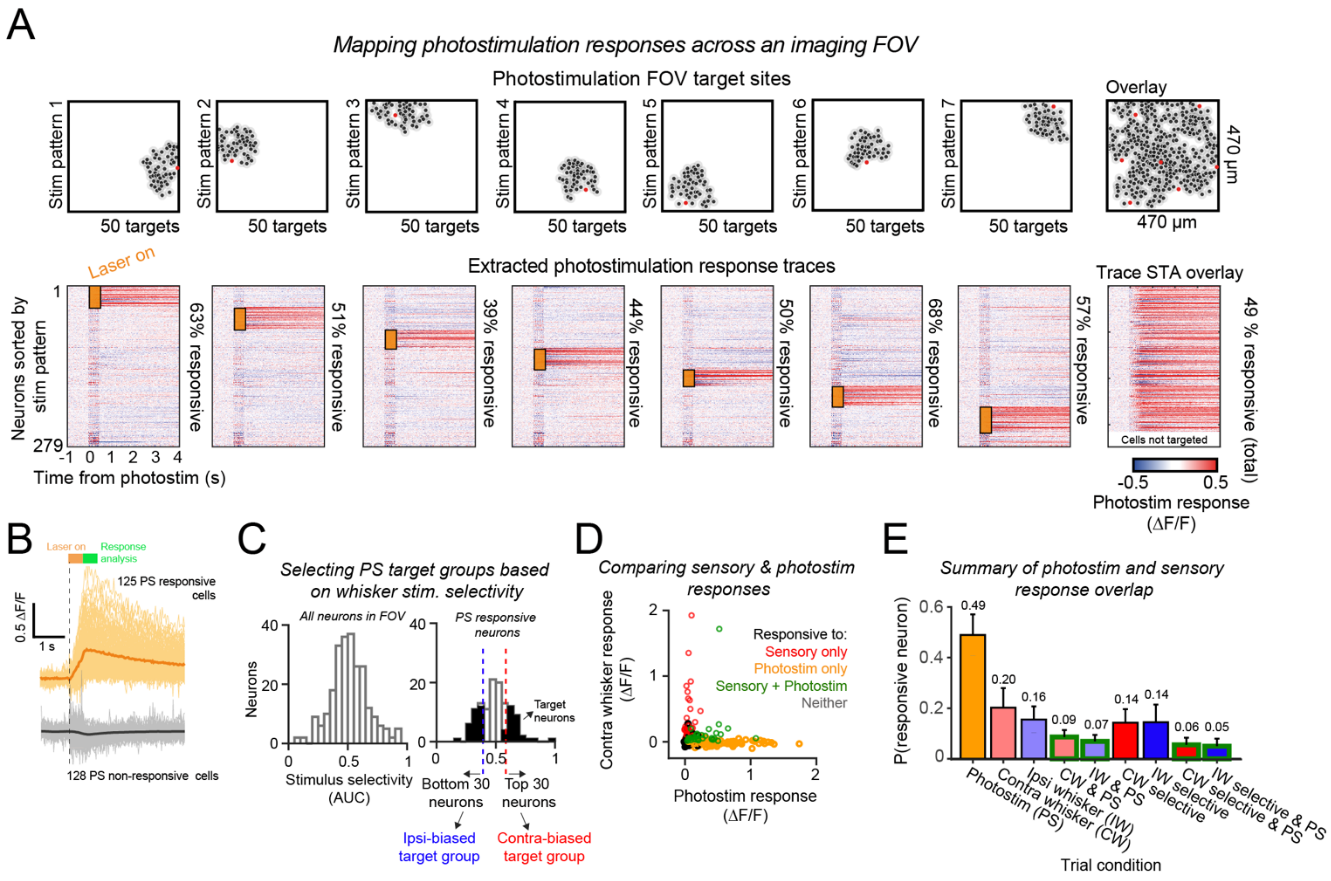


**Figure S9. Selection of two-photon photostimulation target groups**

**(A)** Example photostimulation mapping for an example experiment. All neurons in the FOV were tested for their response to photostimulation. xy pixel locations corresponding to locations of neural somata were detected semi-automatically and were clustered in 7 SLM stimulation patterns with 50 target sites per pattern. Top row: Each column relates to an individual SLM-target pattern within the FOV. The righthand column shows the combined overlay of all patterns. Each target image shows the spiral locations (15 µm diameter; black circle), with a target-zone proximity halo (40 µm diameter; grey circles). Red circles indicate the xy location of the photostimulation galvo mirrors, which were positioned to maximise SLM stimulation efficiency. Bottom row: Average photostimulation-triggered fluorescence traces are shown aligned to stimulation onset (0 s). The orange bars (0 – 0.5 s) indicate the neurons targeted in each stimulation pattern, and the duration of stimulation (500 ms). Neurons are sorted according to stimulation group along the y-axis. Stimulation of each SLM pattern selectively drives activity in the corresponding target neurons. The right image shows the overlay. Note the cells at the bottom of the 'overlay' heatmap were not identified as targets (distance from a spiral location > 20 µm). **(B)** Comparison of PS-evoked fluorescence traces of individual neurons in (A) identified as photostimulation-responsive (orange) and photostimulation non-responsive (black). Each trace shows the average response of a single neuron in (A) when the neuron was targeted for photostimulation. The average across all responsive and non-responsive neuron groups are shown as thicker lines. The green shaded bar indicates the window used to analysis PS responses (500 - 1000 ms post-stimulus onset). **(C)** Overview of target group selection process for an example session. Left: histogram showing whisker stimulus-selectivity scores across all neurons in the FOV. Right: histogram showing stimulus-selectivity scores of PS-responsive neurons in the FOV (as identified in B). Based on this result, photostimulation target groups were designed by selecting the 30 neurons at the top (contra biased) and bottom (ipsi biased) of the distribution. **(D)** Comparison of sensory and photostimulation responses across neurons in an example session. Individual marker points show individual neurons, and the colour denotes whether the neuron was significantly responsive to sensory and/or photostimulation (253 neurons). **(E)** Quantification of the average fraction of neurons significantly responsive to photostimulation and different sensory stimuli across all sessions (52 sessions, 13 mice). Note that finding photostimulation-responsive neurons with significant selectivity for contra vs ipsi whisker input was rare (~5%).

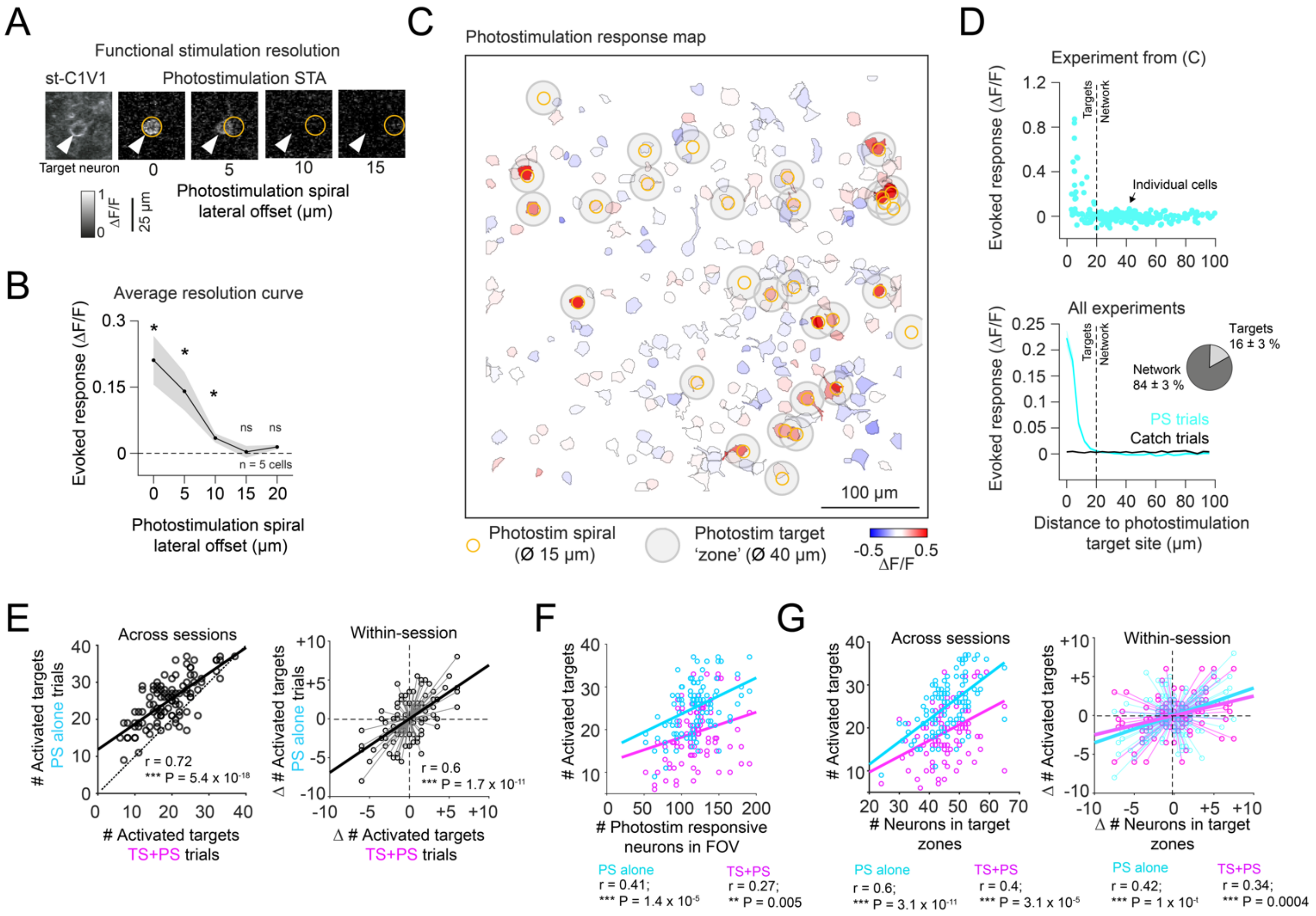


**Figure S10. Functional resolution of targeted photostimulation**

**(A)** Photostimulation resolution was assessed by measuring the fluorescence response (500 - 1000 post-stimulus) in a target neuron as the photostimulation spiral was offset laterally across the cell body. An example pixelwise STA at 0, 5, 10 and 15 µm offset is shown. The white arrow indicates the location of target cell, the orange dashed circle indicates the location of the PS spiral. **(B)** The average PS response of 5 cells as a function of spiral target-site offset as shown in (A). Statistical tests were Wilcoxon signed rank tests difference tested against 0; * P < 0.05. **(C)** Example map showing the photostimulation response for a single SLM-target neuron group (the FOV is the same as in Figure 3D and S1D). The smaller orange circles indicated the locations and size of photostimulation spirals in the FOV, the larger shaded grey circles indicate target 'zones', defined by functional stimulation resolution calibration. Neurons outside target zones ('network' cells), are unlikely to receive direct photo-stimulation. ROIs are segmented by Suite2p and coloured according to mean extracted fluorescence response on PS trials. **(D)** Top: average photostimulation responses extracted from ROIs in the example FOV shown in (C) plotted against distance to nearest photostimulation target site. Each cyan data point represents a single neuron (n = 246 neurons). Bottom: Average photostimulation vs distance to target site responses (binned along the x-axis) for all sessions (cyan). The response on catch trials is shown in dark grey (n = 52 sessions, 2 target-groups per sessions). The vertical dashed line at 20 um shows the target zone threshold. The inset shows the average proportion of target and network cells across all sessions (mean across 104 photostimulation conditions). **(E)** Left: Across session correlation between the number of activated targets on in the presence (x-axis) and absence (y-axis) of sensory (TS) input. Correlation is based on 104 photostimulation conditions (shown as individual markers) in 52 sessions. Right: Correlation between variability in the number of activated targets across the two photostimulation groups (within the same FOV) in the presence (x-axis) and absence (y-axis) of TS input. The data were mean-centred as described in Figure 5E. The correlation coefficient and p-value are indicated on the plot. **(F)** Correlation between the total number of PS-responsive neurons in the FOV (x-axis) and the number of activated targets on PS (cyan) and TS+PS (magenta) trials. **(G)** Right: Comparison between the total number of neurons within photostimulation target zones (x-axis) and the number of activated targets on PS (cyan) and TS+PS (magenta) trials across sessions. Left. Same as Right but showing the comparison across target groups within the same FOV. In all correlation plots, the thick line(s) shows the least-squares fit. The correlation strength (Pearson's corr) and p-value are indicated on the plot. Analyses were based on 104 photostimulation conditions in 52 sessions.

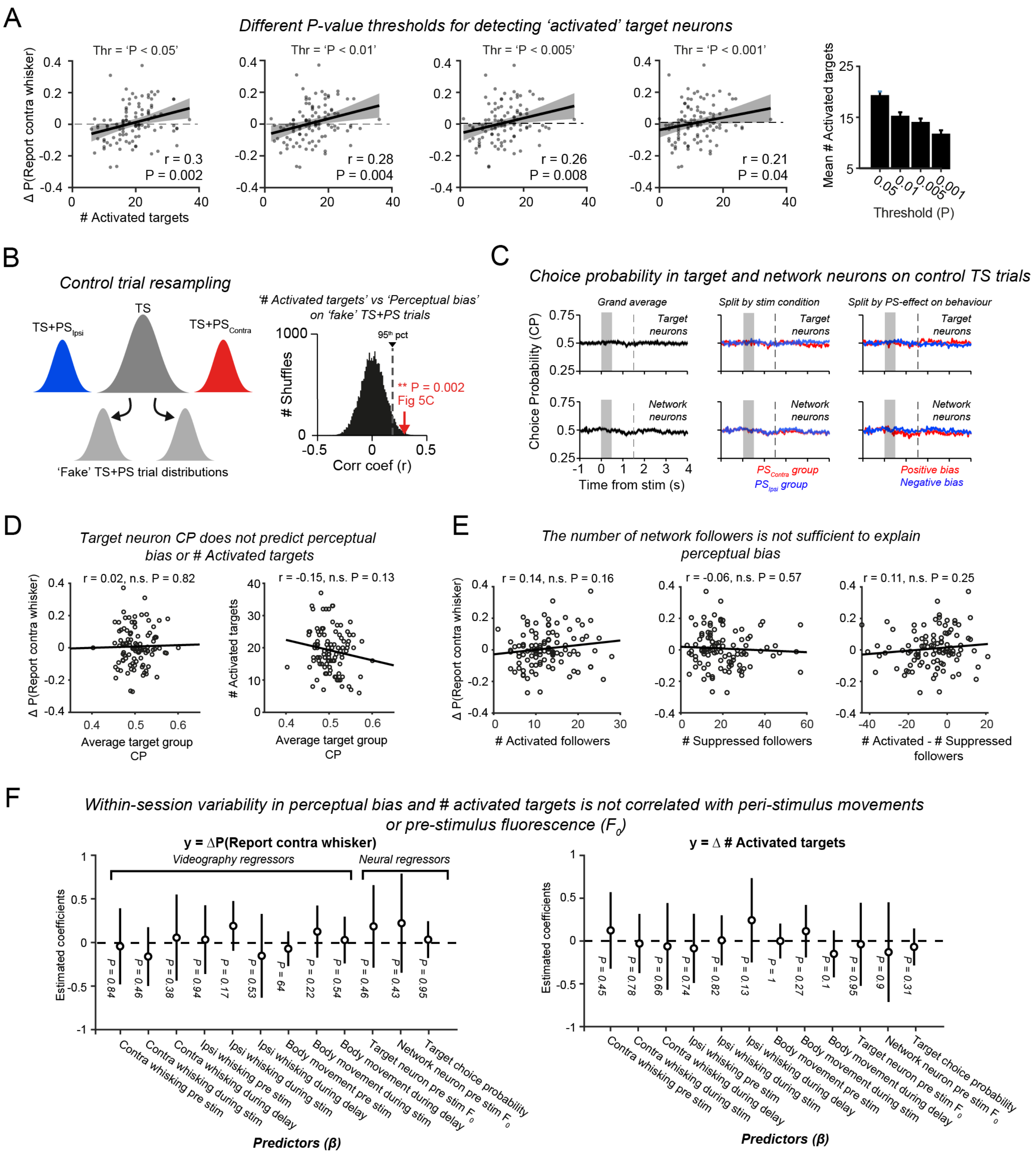
A
Different P-value thresholds for detecting 'activated' target neurons
Thr = 'P < 0.05'
Thr = 'P < 0.01'
Thr = 'P < 0.005'
Thr = 'P < 0.001'
Δ P(Report contra whisker)
### Activated targets
r = 0.3
P = 0.002
r = 0.28
P = 0.004
r = 0.26
P = 0.008
r = 0.21
P = 0.04
Mean # Activated targets
Threshold (P)
B
Control trial resampling
TS
TS+PS Ipsi
TS+PS Contra
'Fake' TS+PS trial distributions
'# Activated targets' vs 'Perceptual bias'
on 'fake' TS+PS trials
### Shuffles
95th pct
** P = 0.002
Fig 5C
Corr coef (r)
C
Choice probability in target and network neurons on control TS trials
Grand average
Split by stim condition
Split by PS-effect on behaviour
Target neurons
Network neurons
Choice Probability (CP)
Time from stim (s)
PS Contra group
PS Ipsi group
Positive bias
Negative bias
D
Target neuron CP does not predict perceptual
bias or # Activated targets
r = 0.02, n.s. P = 0.82
r = -0.15, n.s. P = 0.13
Δ P(Report contra whisker)
### Activated targets
Average target group CP
E
The number of network followers is not sufficient to explain
perceptual bias
r = 0.14, n.s. P = 0.16
r = -0.06, n.s. P = 0.57
r = 0.11, n.s. P = 0.25
### Activated followers
### Suppressed followers
### Activated - # Suppressed followers
F
Within-session variability in perceptual bias and # activated targets is not correlated with peri-stimulus movements
or pre-stimulus fluorescence (F0)
y = ΔP(Report contra whisker)
y = Δ # Activated targets
Videography regressors
Neural regressors
Estimated coefficients
P = 0.84
P = 0.46
P = 0.38
P = 0.94
P = 0.17
P = 0.53
P = 64
P = 0.22
P = 0.54
P = 0.46
P = 0.43
P = 0.95
P = 0.45
P = 0.78
P = 0.66
P = 0.74
P = 0.82
P = 0.13
P = 1
P = 0.27
P = 0.1
P = 0.95
P = 0.9
P = 0.31
Contra whisking pre stim
Contra whisking during stim
Contra whisking during delay
Ipsi whisking pre stim
Ipsi whisking during stim
Ipsi whisking during delay
Body movement pre stim
Body movement during stim
Body movement during delay
Target neuron pre stim F0
Network neuron pre stim F0
Target choice probability
Predictors (β)

**Figure S11. The correlation between activated targets and perceptual bias is not explained by other neural and behavioural factors.**

**(A)** Correlation plots showing perceptual bias (y-axis) against the number of activated target neurons (x-axis) when using different p-value thresholds for detecting activated targets. The correlation strength and significance is indicated on each plot. The plot on the far right shows the average number of neurons identified as significantly activated as a function of p-value thresholds. Correlations are based on 104 photostimulation conditions in 52 sessions. Shaded error bars show 95% confidence intervals of the linear regression fit. **(B)** Left: Schematic overview of the procedure for resampling TS control trials. Right: Distribution of the correlation coefficients (Pearson’s corr r) obtained across 10000 iterations of the shuffled resampling procedure. The probability of obtaining a correlation r = 0.3 (as in Figure 5C) across sessions between target activation and perceptual bias on control trials was 0.002 (indicated by the red arrow). **(C)** Average choice probability in target groups (top row) and network neuron groups (bottom row) across sessions. The left column shows the grand average, the middle column shows groups split by contra vs ipsi-biased target group, and the right column shows target groups split by positive vs negative perceptual bias effect. **(D)** Target group choice probability does not correlate with perceptual bias (left), or with the number of activated target group neurons (right). The correlation strength (Pearson’s corr) and significance is indicated on each plot. **(E)** The number of activated (left), suppressed (middle) or difference between activated and suppressed (right) followers does not predict the perceptual effect of PS on choice bias. **(F)** Summary results of a multiple linear regression analysis testing the relationship between within-session variability in perceptual bias (left) and within-session variability in target activation (right) and differences in number of behavioural and neural predictors measured across the two photostimulation groups. Behavioural predictors included quantification of pre (-1000 to 0 ms), during (0 to 500 ms) and post (500 to 1000 ms) stimulus whisking and body movement. Neural predictors included average pre-stimulus (-1000 to 0 ms) baseline $F_0$ fluorescence in target and network neurons groups and target group choice probability. Marker points show the estimated coefficients and 95% confidence intervals, with the p-values denoting significance of each predictor. Data in this figure come from n = 104 target groups across 52 sessions; 13 mice.

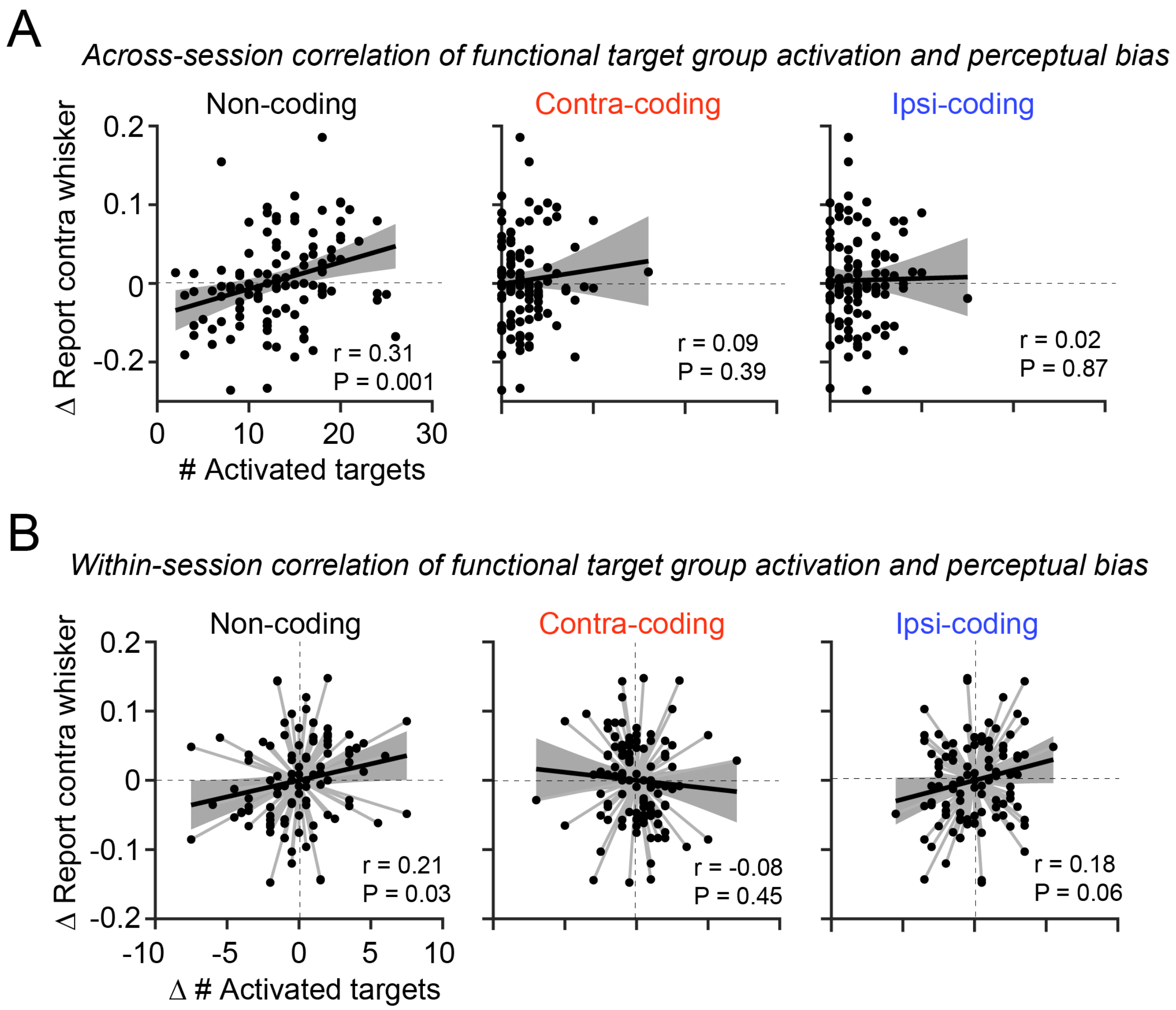


**Figure S12. The number of non-coding targets activated predicts perceptual bias across and within sessions.**

**(A)** Perceptual bias (y-axis) is plotted against the number of activated targets (x-axis) in non-coding (left), contra-coding (middle) and ipsi-coding (right) categorical groups across sessions. The line and shaded error bars show the regression fit and 95% confidence intervals. The correlation strength (Pearson's corr) and significance are shown in each plot. **(B)** Same as (A) but showing the within-session comparison between photostimulation groups. Data come from 104 photostimulation conditions in 52 sessions. Each marker indicates an individual photostimulation condition.

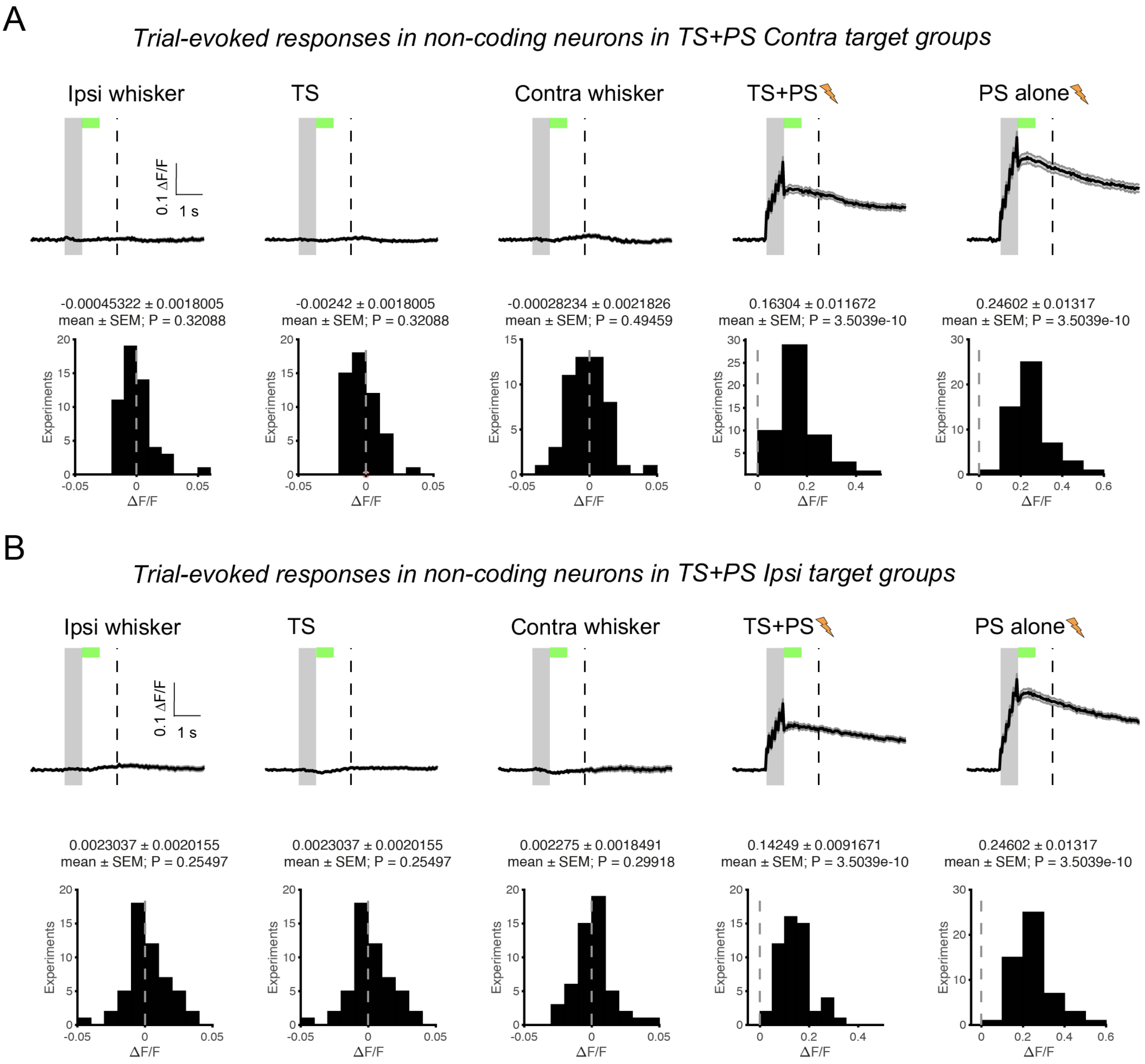


**Figure S13. Trial-evoked activity in non-coding target neurons split by photostimulation target group.**

**(A)** Top row: Average evoked fluorescence traces in non-coding target neurons within the contra-biased PS target group across trial-types (as shown in Figure 6). The grey shaded bar shows the stimulus presentation period, and the vertical dashed line shows the time of the go cue. Bottom row: Histogram of average evoked response in the non-coding target group (quantified 500 - 1000 ms post stimulus; green shaded bar in top row) is shown for all sessions (n = 52 sessions; 13 mice). The average response across all sessions in stated in the header text of each plot and the P-value shows the result of a Wilcoxon signed rank test comparing the distribution to 0. **(B)** same as in (A) but showing quantification of responses in non-coding neurons within the ipsi-biased PS target groups.
